# Establishing Probiotic *Saccharomyces boulardii* as a Model Organism for Synthesis and Delivery of Biomolecules

**DOI:** 10.1101/2020.01.22.915389

**Authors:** Deniz Durmusoglu, Ibrahim Al’Abri, Scott P. Collins, Chase Beisel, Nathan Crook

**Author notes:** These authors contributed equally to this work.

## Abstract

*Saccharomyces boulardii* is a widely used yeast probiotic which can counteract various gastrointestinal disorders^1^. As a relative of *Saccharomyces cerevisiae*, *S. boulardii* exhibits rapid growth and is easy to transform^2^ and thus represents a promising chassis for the engineered secretion of biomolecules. To establish *S. boulardii* as a platform for delivery of biomolecules to the mammalian gut, we measured the amount and variance in protein expression enabled by promoters, terminators, selective markers, and copy number control elements in this organism. These genetic elements were characterized in plasmidic and genomic contexts, revealing strategies for tunable control of gene expression and CRISPR-mediated genome editing in this strain. We then leveraged this set of genetic parts to combinatorially assemble pathways enabling a wide range of drug and vitamin titers. Finally, we measured *S. boulardii*’s residence time in the gastrointestinal tracts of germ-free and antibiotic-treated mice, revealing the relationships between dosing strategy and colonization level. This work establishes *S. boulardii* as a genetically tractable commensal fungus and provides a set of strategies for engineering *S. boulardii* to synthesize and deliver biomolecules during gut colonization.

## Introduction

Next-generation probiotics offer unique attributes for the delivery of biomolecules to the gut^3^. By leveraging decades of work in metabolic engineering^4^, engineered probiotics have enabled the conversion of unused dietary material to beneficial products for the host^5–7^. Furthermore, the ongoing development of synthetic biological sensors advance the concept of drug synthesis tailored to the severity and location of disease within the host^8–11^. Coupled with existing manufacturing expertise in food science and infrastructure in place for distributing probiotics, engineered probiotics promise to substantially reduce the costs associated with production of drug molecules^12^. All engineered probiotics described to date have been members of the bacterial domain of life due to their high numerical abundance in the gut^13, 14^ and ease of engineering.^9, 15^ For example, *Escherichia coli* Nissle 1917 has been engineered to treat diseases such as phenylketonuria (PKU)^15, 16^, hyperammonemia^17^, and cancer^18^, or eliminate pathogens such as *Pseudomonas aeruginosa*^19^. In addition to these advantages, the limitations to bacterial probiotics include susceptibility to antibiotics,^20^ predation by bacteriophage known to be highly abundant in the human gut,^21^ and difficulty in producing high levels of post-translationally-modified proteins^22^.

In addition to bacteria, a diverse fungal population also exists in the human gastrointestinal tract (GIT)^23–26^. While the numerical abundance of fungal cells in the GIT is much lower than bacterial cells, fungal cells are on average 100-fold larger, indicating that their biomass and role in the gut may be larger than metagenomic surveys indicate^25, 27^. Recent studies have shown that while commensal fungi play important roles in the development of inflammatory bowel diseases, they also provide protective heterologous immunity to pathogenic organisms by training the immune system^28, 29^. Furthermore, fungi are not susceptible to bacteriophage predation and are easily engineered to secrete high titers of proteins that are post-translationally modified. Thus, in this study, we established a pipeline to engineer the only eukaryotic probiotic that is approved by the FDA: *Saccharomyces boulardii*^30^.

*S. boulardii* is a non-pathogenic yeast that was originally isolated from lychee and mangosteen in 1923 and has since been used to treat ulcerative colitis, diarrhea, and recurrent *Clostridium difficile* infection^1, 31^*. S. boulardii* is closely related to the famous budding yeast *Saccharomyces cerevisiae*, indicating that it may be similarly amenable to engineering for the production of biomolecules, including biologics requiring post-translational modifications^2, 32^. Supporting this, *S. cerevisiae* expression vectors can be transformed and propagated in *S. boulardii*, and CRISPR/Cas9-mediated genome editing is functional in both strains^33^. However, compared to *S. cerevisiae, S. boulardii* better tolerates low pH environments and grows more rapidly at human body temperature^32, 34, 35^. Given its regulatory status, genetically tractability, and well-studied yeast relative, *S. boulardii* is uniquely poised as a potential model chassis for the delivery of therapeutics to the human gut. Indeed, *S. boulardii* has been engineered to secrete human synthetic lysozyme^33^ and HIV-1 Gag^36^ in cell culture, as well as IL-10 in an IBD mouse model^37^. However, while the rules governing high-level biomolecule production in *S. cerevisiae* have been extensively studied, these rules remain to be defined in culture- and gut-resident *S. boulardii*. In this work, we sought to develop a quantitative framework for engineering *S. boulardii* for *in situ* production and delivery of therapeutics to the mammalian gut. We began by assaying the ability of a library of *S. cerevisiae* genetic parts^38^ to drive gene expression from various plasmidic and genomic contexts in *S. boulardii*. We then applied these genetic parts to rapidly construct and identify *S. boulardii* strains exhibiting high levels of vitamin (β-carotene) or drug (violacein) production. Finally, we measured the residence time of *S. boulardii* in the mouse gut. Taken together, this work demonstrates the promise of, and details a set of procedures for engineering, *S. boulardii* for producing biomolecules in the gut.

## Results

### Selective marker and plasmid origin tune gene expression over a wide range in *S. boulardii*

Plasmids enable rapid prototyping of synthetic constructs in many organisms, including yeast. The tunability of gene expression on plasmids is due in part to plasmid-specific copy number control sequences, which in yeast have been shown to include the origin of replication (origin) and selectable marker (marker)^39^. We therefore first sought to understand the interplay between origin and marker in determining the expression of a model fluorescent reporter gene. An optimal reporter gene, when expressed, should yield detection signals well above the background signal produced by non-expressing strains in order to precisely resolve gene expression levels^40^. Therefore, we first measured the cell fluorescence conferred by three proteins (*Venus, mTurquoise2, mRuby2*) when expressed by the *TDH3* promoter and the *TDH1* terminator on a plasmid harboring a *URA3* marker and the 2μ origin. Fluorescence levels were measured at log phase using flow cytometry. In all experiments, we normalized the resulting fluorescence levels to those of a strain harboring an empty vector. *Venus* displayed the highest normalized fluorescence (1080-fold) (Supplementary Figure S1). However, *Venus* fluorescence also exhibited high cell-to-cell variability compared to that for *mRuby2* and *mTurquoise2*, and its fluorescence distribution was non-gaussian, suggesting that *Venus* would not be suitable for measuring fine differences in expression. On the other hand, *mRuby2* showed the second highest normalized fluorescence (450-fold) and exhibited a more gaussian fluorescence distribution. Because *mRuby2* exhibited a 2.6-fold higher normalized fluorescence than *mTurquoise2*, we decided to proceed with *mRuby2* for further analysis of synthetic construct performance in *S. boulardii*.

We generated expression cassettes containing the *TDH3* promoter (*pTDH3*), *mRuby2*, and the *TDH1* terminator (*tTDH1*) in combination with 3 auxotrophic markers (*URA3*, *HIS3* and *TRP1*) or four antifungal markers (geneticin resistance (*KanMX*), zeocin resistance (*ZeoR*), nourseothricin resistance (*NatR*), and hygromycin resistance (*HygR*)) and two origins-of-replication (CEN6/ARS4 (*CEN*, low-copy) and 2-micron (2μ, high-copy)). We transformed the cassettes containing auxotrophic markers into corresponding auxotrophic strains (*S.bΔURA3*, *S.bΔHIS3 and S.bΔTRP1)* and antifungal markers into wild-type strain. We achieved a maximum normalized fluorescence of 400 using *URA3* and 2μ (Figure 1a, Supplementary Figure S2a). Among antifungal markers, *HygR* coupled with the 2μ origin yielded the highest fluorescence (Figure 1a). Other marker/origin combinations yielded a 100-fold variation in fluorescence (Figure 1a, Supplementary Figure S2). This large range in expression levels for constructs containing the same promoter and terminator indicated that, as in *S. cerevisiae*, plasmid origin and marker have significant effects on gene expression in *S. boulardii*.

**Figure 1.**
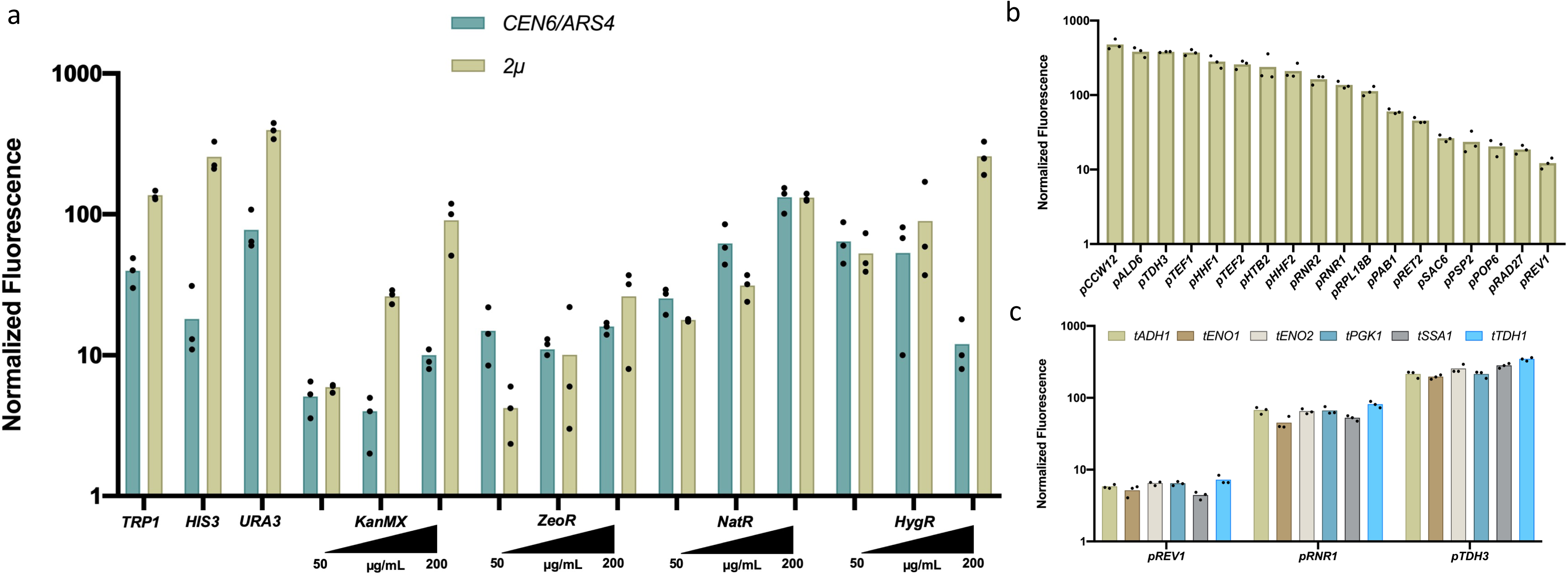
Characterization of *S. boulardii* (a) markers and origins of replication, (b) promoters, and (c) terminators *in vitro.* **a)** 3 auxotrophic (*URA3, HIS3, TRP1*) and 4 antifungal (*KanMX, ZeoR, NatR, HygR*) markers were cloned in plasmids containing the *TDH3* promoter, mRuby2 fluorescent protein and *TDH1* terminator. Centromeric (*CEN6/ARS4*) and episomal (*2µ*) origins were cloned in combination with the 7 selective markers. Plasmids with antifungal markers were tested in 3 different antifungal concentrations (50, 100 and 200 µg/mL). **b)** 18 constitutive promoters were cloned 5’ of the *mRuby2* fluorescent protein and the *TDH1* terminator. **c)** 5 terminators were cloned 3’ of 3 different promoters at different strength (*pTDH3*-strong, *pRNR1*-medium and *pREV1*-weak) and the *mRuby2* fluorescent protein. Absolute fluorescence values were obtained via flow cytometry for 3 biological replicates (n=3). Absolute fluorescence values were normalized to fluorescent values of cells harboring the spacer plasmid. The height of the bars represents the normalized mean fluorescent value of 3 biological replicates and the black dots show the normalized mean of each biological replicate.

### The impact of origin on gene expression is marker-dependent in *S. boulardii*

We next asked whether certain markers or origins had consistent effects on fluorescence. For auxotrophic markers, 2μ plasmids consistently yielded higher fluorescence than CEN plasmids (Two-way ANOVA, p<0.05) (Figure 1a). This is consistent with prior work in *S. cerevisiae*, which demonstrated that 2μ plasmids with auxotrophic markers are maintained at 28-58 copies per cell, whereas CEN plasmids are maintained at only 4-8 copies per cell^39^. However, with the exception of *HygR* (Two-way ANOVA, p<0.0001), the fluorescence conferred by plasmids containing antifungal markers was not impacted by the choice of origin (Two-way ANOVA, p>0.05). For the CEN origin, using *NatR* at the maximum antifungal concentration yielded the highest fluorescence level. This level was significantly higher expression than those conferred by the *HIS3*, *ZeoR*, *KanMX*, or *HygR* markers (Two-way ANOVA, p<0.05). However, for the 2μ origin, using any of the auxotrophic markers gave gene expression levels higher than any antifungal marker (Two-way ANOVA, p<0.0001). Taken together, our data indicate that gene expression conferred by plasmids containing auxotrophic markers (but not antifungal markers) is sensitive to the choice of origin, and that selective marker significantly impacts gene expression for both 2μ and CEN origins.

### Plasmidic gene expression noise is marker and origin dependent in *S. boulardii*

Cell-to-cell variability in gene expression can have dramatic impacts on overall production titers^41^. Therefore, it is desirable to design engineered probiotics to exhibit minimal gene expression noise. Previous work characterizing gene expression noise in *S. cerevisiae* showed that cell variability varies as a function of promoter, terminator, origin, and selective marker^42^. We thus calculated the population-level coefficient of variance (CV) in the fluorescence values we obtained for *S. boulardii*. We found that marker, origin, and marker-origin interactions had significant impacts on expression noise (Two-way ANOVA, p<0.05). Specifically, CEN plasmids exhibited less cell-cell variability than 2µ plasmids (Two-way ANOVA, p<0.05). Additionally, antifungal markers exhibited less noisy expression than auxotrophic markers (Two-way ANOVA, p<0.05). These data suggest differences in the ways that the auxotrophic and antifungal markers interact with origins of replication to control gene expression, and point to design strategies which reduce plasmid-based gene expression variance.

*S. cerevisiae* promoters largely maintain their relative activities in *S. boulardii*. Promoter sequences are frequently employed to tune the expression of individual genes in yeast^43, 44^. Here, we quantified the transcriptional strength of 18 constitutive *S. cerevisiae* promoters driving the expression of a fluorescent protein in *S. boulardii*. Each of these promoters had a homolog in *S. boulardii*, with percent nucleotide identities ranging from 100% to 97% (Supplementary Table S1). In these experiments, vectors contained the 2μ origin and the *URA3* auxotrophic marker, which maximized the range over which changes to fluorescence could be detected (Figure 1a). Each promoter drove the expression of *mRuby2* and transcription was terminated by *tTHD1*. These constructs were transformed into *S. boulardii* and fluorescence levels were measured at log phase via flow cytometry. Each promoter yielded different mean fluorescence values (Two-way ANOVA, p<0.0001), and these promoters collectively modulated fluorescence levels over a 40-fold range (Figures 1b, S3). While the relative activities of 17 of the 18 promoters in *S. boulardii* were similar to reported values in *S. cerevisiae*, one promoter (*pALD6*) showed 10-fold higher activity in *S. boulardii* than in *S. cerevisiae* (Supplementary Figure S4). While the precise role of *ALD6* in the probiotic properties of *S. boulardii* is unknown, Ald6p is involved in the detoxification of aldehyde molecules to their corresponding acid in *S. cerevisiae*^45^. Collectively, this data curates a set of promoters enabling precise tuning of gene expression in *S. boulardii*, and indicates that constructs developed for *S. cerevisiae* will exhibit similar gene expression profiles in *S. boulardii*.

### Terminators modulate gene expression in a promoter-dependent manner in *S. boulardii*

Terminators regulate expression in yeast by modulating mRNA stability, translational efficiency and mRNA localization^46^. Furthermore, a terminator’s effect on expression can depend on the strength of the promoter or the sequence of the expressed gene^47, 48^. Therefore, we were interested in defining the effect of terminator choice on gene expression in *S. boulardii*. We cloned 6 terminators (*tENO1, tENO2, tSSA1, tPGK1, tADH1,* and *tTDH1*) into mRuby2 expression constructs driven by strong (*pTDH1*), medium (*pRNR1*), or weak (*pREV1*) promoters. We found that these terminators could modulate gene expression by 1.77, 1.81, and 1.64-fold in constructs containing *pTDH1*, *pRNR1*, or *pREV1*, respectively, effectively increasing the expression range beyond that enabled by varying the promoter alone (40-fold, Figure 1b). In particular, using a strong promoter (*TDH3*) with the *TDH1* terminator yielded 80-fold higher expression than a weak promoter (*REV1*) with the *SSA1* terminator. Relative to *tSSA1 and tTDH1,* we found that *tENO1*, *tENO2, tADH1* and *tPGK1* significantly decreased gene expression in constructs containing the strong *pTDH3* promoter (Two-way ANOVA, p<0.005) while for *pRNR1* and *pREV1*, the differences in gene expression we observed among terminators were not statistically significant (Two-way ANOVA, p>0.05) (Figure 1c, S5). Taken together, this data suggests that varying the terminator is a valuable way to fine-tune gene expression, particularly when certain regulatory characteristics of the promoter (e.g. responsiveness to growth phase or gut conditions) must be maintained.

### The *S. boulardii* genome can be efficiently modified by CRISPR-Cas genome editing

Genomic integration of synthetic constructs can sidestep issues associated with the use of plasmids in the gut. Such issues include plasmid instability if no selective pressure is present, potential spread to other microbes, collateral damage to the microbiota if antimicrobials are used to select for plasmid maintenance, and the increased metabolic burden associated with the maintenance of multicopy plasmids. However, different genomic regions are known to support varying levels of synthetic construct expression, potentially due to nucleosome positioning^49^, and integrating material into the genome can be especially difficult for diploid and polyploid organisms^50^.

To characterize and overcome inefficiencies associated with genome integration, we compared three different editing methods in *S. boulardii*: un-assisted linear dsDNA integration (dsDNA integration) and genome editing assisted with the CRISPR nucleases SpCas9 or LbCas12a . We tested the ability of these methods to integrate a linear expression cassette containing *mTurquoise2* in three different loci (INT1, 4 and 5) via colony PCR. In the case of dsDNA integration, 4/12 transformants did not contain the expected *mTurquoise2* insertion, while 7/12 colonies yielded multiple PCR fragments (Supplementary Figure S6). Sequencing revealed that the larger fragment corresponded to the desired integration, while the shorter fragment corresponded to the native genomic locus. This result is likely due to the fact that the *S. boulardii* genome is diploid, and that integration of one copy of the construct is sufficient to confer uracil prototrophy.. Therefore, we tested whether CRISPR-based editing tools could achieve more complete genome editing in *S. boulardii*. We reasoned that since CRISPR-Cas systems enhance the proportion of double-stranded breaks in all copies of the targeted sequence, editing of all chromosomal copies would be a preferred survival strategy. Three sgRNAs for SpCas9 and three gRNAs for LbCas12a were designed for each integration site. Targeted loci were followed by a 5’-NGG-3’ protospacer adjacent motif (PAM) for SpCas9 and preceded by a 5’-TTTV-3’ PAM (V = A, C, G) for LbCas12a. Supplementary Figure S7c-f shows the plasmid designs we used for CRISPR-based editing, Supplementary Table S2 shows integration locations (INT1-INT5) in *S. boulardii*’s genome, and Supplementary Table S3 shows the guide RNA sequences we used.

We tested the editing efficiency at each integration site (using a single guide RNA per experiment) as above for un-assisted linear DNA integration. Supplementary Table S4 shows the number of fluorescent colonies we observed for each editing method. Colony PCR of 4 random fluorescent transformants from each experiment revealed a single band corresponding to either the edited or the unedited version of the locus (Supplementary Figure S8). Altogether, we were able to generate genome integrants with 82-88% efficiency for SpCas9 and 93-99% efficiency for LbCas12a in *S. boulardii* (Figure 2a, Supplementary Table S4). Thus, these results indicate that CRISPR/LbCas12a-assisted genome editing has >10% and >50% higher editing efficiency than CRISPR/SpCas9 and dsDNA integration, respectively, and is therefore a promising choice for future editing in this strain.

**Figure 2.**
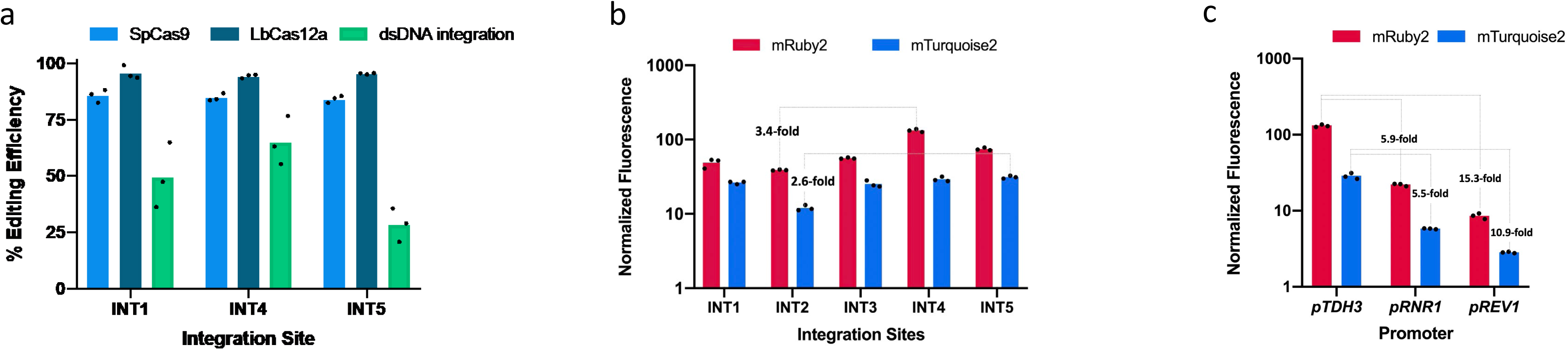
*S. boulardii* genomic location impacts gene expression, and comparison of genome editing techniques. **a**. Editing efficiency at 3 different loci using 3 different methods: CRISPR/SpCas9 (SpCas9), CRISPR/LbCas12a (LbCas12a) and un-assisted linear dsDNA integration (dsDNA integration). 3 different guides were used for both SpCas9 and LbCas12a for each site. Each dot represents one guide. 3 replicate cultures were transformed with the DNA repair template to measure the efficiency of genome editing in *Sb* via dsDNA integration. Table S1 shows the number of colonies screen for each method. **b.** Effect of chromosomal locus on the expression of the fluorescent genes mRuby2 and mTurqouise2. The dotted lines show the fold change between the expression of mRuby2 and mTurquoise2 between the highest and lowest expression among all the integration sites. Each dot represents one biological replicate. **c.** The effects of promoter (strong, medium, or weak) on gene expression at the same chromosomal locus. The dotted lines show the fold change between the expression of *mRuby2* and mTurquoise2 using different promoters. Each dot represents one biological replicate.

### Integration locus has a minor effect on gene expression, relative to promoter choice

To determine the extent of locus-dependent effects on gene expression in *S. boulardii*, we integrated constructs driving *mRuby2* or *mTurquoise2* expression into five different locations in the *S. boulardii* genome (Supplementary Table S2) using LbCas12a. These genes were regulated by *pTDH3* and *tTDH1*, and constructs contained *URA3* to select for edited strains (Supplementary Figure S7b). The *mRuby2* and *mTurquoise2* fluorescent proteins are sequence-divergent, with only 30% amino acid identity. These two proteins therefore enabled a cursory view into the generalizability of these patterns across genes. Across integration sites, the expression of *mRuby2* and *mTurquoise2* varied 3.4-fold and 2.6-fold, respectively (Figure 2b, S9). INT2 supported the lowest expression for both genes, while INT4 supported the highest expression levels for *mRuby2* and INT5 supported the highest expression level for *mTurquoise2*. We next asked whether differences in promoter strength observed in the plasmid-based system would be similar for genomically-integrated constructs. We therefore expressed *mRuby2* and *mTurquoise2* from two additional promoters in INT4: *pRNR1* and *pREV1*. Intriguingly, genomic *mRuby2* expression varied by 5.9-fold between *pTDH3* and *pRNR1*, compared with 2.8-fold for the same construct on a 2µ plasmid. Conversely, genomic *mRuby2* expression varied by 15.3-fold between *pTDH3* and *pREV1*, compared to 31.2-fold for the same promoters used on a 2µ plasmid (Figure 2c). This could indicate that gene expression from high-copy plasmids saturates at high promoter strengths, or that promoter strength varies with copy number due to transcription factor dilution. Together, this work establishes a pipeline for efficient genome integration in *S. boulardii* and identifies regions of the *S. boulardii* genome enabling high-level expression of synthetic constructs. Further, these data show that the impact of integration locus on gene expression is relatively small in *S. boulardii* compared to that of the promoter.

### *S. boulardii* facilitates high-throughput metabolic engineering through *in vivo* combinatorial assembly of multi-enzyme pathways

Having found that *S. boulardii* could support genome integration via homologous recombination, we asked if recombination efficiencies were sufficient for *in vivo* assembly of biosynthetic pathways. In *S. cerevisiae,* combinatorial *in vivo* assembly is a facile technique for rapid optimization of metabolic pathways^51–53^. Large scale assembly of common heterologous metabolic pathways directly in *S. boulardii* would simplify strain development and might reveal the potential for *S. boulardii* as a host for biomolecule production.

We implemented golden gate assembly to streamline *in vivo* assembly workflows in *S. boulardii*. Namely, we started from our characterized promoter, coding sequence (CDS), and terminator parts and assembled pathways directly in yeast with minimal subcloning (Figure 3a). Our assembly strategy involved two steps: a golden gate assembly step to assemble a 9-member gene-level library of promoter-CDS-terminator transcriptional units for each gene in the pathway of interest and *in vivo* homologous recombination to assemble the gene-level libraries created in step 1 into pathways, as shown in Figure 3a. Each gene-level library member differed only in the promoter driving gene expression. These transcriptional units are flanked with homology arms which facilitate *in vivo* assembly. These gene-level libraries are then PCR-amplified directly from the Golden Gate reaction mixture. Purified amplicons are transformed into *S. boulardii,* where flanking sequences direct homology-dependent assembly of the final pathway with one variant of each transcriptional unit. We used this method to construct β-carotene and violacein synthesis pathways, which provided a direct colorimetric readout of assembly efficacy (Supplementary Figures S10a and S10b). Supplementary Figures S10c and S10d show spotted cultures of 28 library members from the β-carotene pathway and 24 library members from the violacein pathway. Colony sequencing revealed that each strain harbored a plasmid containing the expected pathway topology, with each gene driven by one of the 10 selected promoters and the desired terminator.

**Figure 3.**
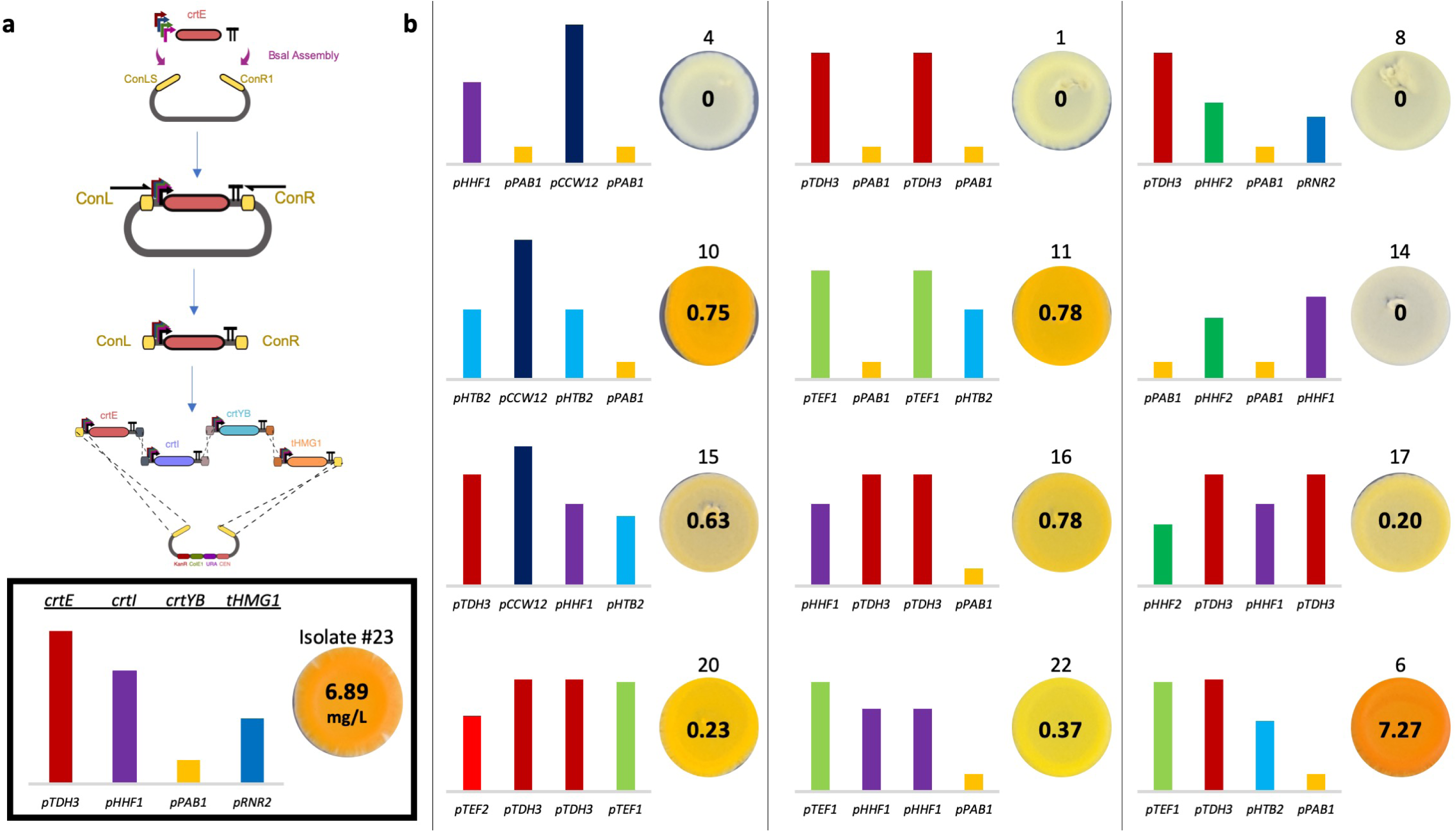
Combinatorial Assembly of β-carotene Pathway in *Saccharomyces boulardii*. **a)** Overview of assembly steps. Genes in the β-carotene pathway were randomly cloned behind 9 promoters (*pTDH3, pCCW12, pHHF2, pTEF1, pTEF2, pHHF1, pHTB2, pPAB1, pRNR2). tSSA1, tADH1, tTDH1 and tENO1* terminators were cloned behind *crtE, crtI, crttYB-2* and *tHMG1*, respectively. The backbone plasmid contains Type1 and Type2 YTK connectors to enable in vivo *S. boulardii* assembly. Promoter-gene-terminator regions between the two connectors were amplified and the PCR products were transformed in *S. boulardii* for pathway assembly into a linear backbone via connector-assisted homologous recombination, as described in the Methods Section. **b)** Sequencing and β-carotene quantification results of isolates from yeast transformation plates (**Supplementary Figure S8b**). 13 out of 28 isolates (Supplementary Figure S8d) were selected to proceed with sequencing and β-carotene production quantification based on stability of colony color after 3 serial passages. Promoter sequences were confirmed by Sanger sequencing with reverse primer binding on the gene of interest (backbone for *tHMG1*) and forward primer binding on the terminator of the previous gene (backbone for *crtE*). Quantification of β-carotene was done by high-performance liquid chromatography (HPLC) as described in the Methods section. The spot images were taken from Supplementary Figure S8d. Number above the spots corresponds to the spot number on isolate plate on Supplementary Figure S8d. The bar heights represent the normalized fluorescence values in linear scale for each promoter according to promoter characterization work (Figure 1a). The numbers in the center of the spots corresponds to average of β-carotene produced by 3 biological replicate of each isolate in one liter of saturated yeast culture.

We next focused on those strains exhibiting qualitatively stable pigment levels over three rounds of growth of solid media. 13 out of 28 strains and 8 out of 24 strains stably produced β-carotene and violacein molecules, respectively. We determined the identity of the promoters driving each gene in these strains. We also quantified the amount of β-carotene produced by these strains to understand how different combinations of promoters regulate biomolecule production. Figures 3b and S11 show the strength of the promoters comprising each assembled pathway and amount of β-carotene or violacein produced by each assembled pathway. Interestingly, we found that strains with the highest qualitative color intensity and quantitative β-carotene concentration did not harbor strong promoters for all the genes in the pathway. In fact, many high producers contained moderate or low-strength promoters in at least one of their genes. Additionally, we observed cases where multiple pathway genes in a high-producing strain were driven by the same promoter. This suggests that product titers in these strains were not sufficiently high to select for the emergence of non-producers via homologous recombination.

The optimal expression levels for several genes in the β-carotene are not obvious from the biosynthetic reaction map^54^. *HMG1* diverts flux from cellular acetyl-coA pools into the β-carotene pathway, thereby reducing the amount of acetyl-coA available for other processes. *crtYB* can convert lycopene into the desired product β-carotene, but can also convert neurosporene (a precursor to lycopene) into the side product 7,8-dihydro-β-carotene (β-zeacarotene). *crtI* operates on several pathway intermediates, including phytoene, ζ-carotene, and neurosporene^55^. Figure 3b shows substantial variations in β-carotene titer depending on the expression level of these genes. As complex tradeoffs between gene expression and pathway output will likely also be present in engineered strains colonizing the gut, combinatorial assembly is useful for rapidly generating strain designs for subsequent *in vivo* testing or screening. Our results reveal that high biomolecule titers can be achieved via expression tuning, confirming previous results from other methods for pathway optimization in *S. cerevisiae*^52, 53, 56^. Moreover, as in *S. cerevisiae*, large combinatorial pathway libraries can be easily generated in *S. boulardii*, enabling rapid optimization of biomolecule titers.

### The Mouse Microbiota Restricts *S. boulardii* Colonization

Host immunity, intestinal peristalsis, inter-microbial competition, and nutrient availability shape the residence time of microbes in the gut. To parse this complexity, gnotobiotic mouse models are attractive to explore the behavior of microbes in defined gut ecologies. Thus, we first measured *S. boulardii’*s residence time in the germ-free gut by delivering *S. boulardii* (10^8^, 10^7^, or 10^6^ colony-forming units) to germ-free mice via oral gavage (Supplementary Figure S12). No deleterious effects on mouse health were apparent during the experiment. While all dosages yielded similar fecal loads of *S. boulardii* through day 5 (3x10^6^ CFU/g feces), we found that when mice were provided at least 10^8^ CFUs, consistent fecal loads of 10^7^ CFU/g *S. boulardii* were maintained for over 30 days. On the other hand, mice gavaged with 10^6^ and 10^7^ CFUs showed 3-fold lower fecal titers starting on day 7. Thus, the initial dose of *S. boulardii* impacts its fecal titer and residence time in germ-free mice. Furthermore, in the absence of microbial competitors, *S. boulardii* is capable of long-term (>30 days) colonization of the GIT.

We next asked whether the presence of microbial competitors would modulate the gut residence time of *S. boulardii*. To answer this, we turned to conventionally-raised mice treated with antibiotics^57^. In these experiments, we were initially unable to track *S. boulardii* titers via plating because other native gut fungi outgrew *S. boulardii* on YPD plates containing 0.125 mg/ml penicillin and 0.25 mg/ml streptomycin. Therefore, we hypothesized that integration of an antifungal resistance gene into the *S. boulardii* genome would allow easier identification via plating on media containing antifungals. Based on the observed minimum inhibitory concentrations (MICs) of various antifungals against *S. boulardii* (Supplementary Figure S13), we integrated the nourseothricin resistance gene (*NatR*) into the *S. boulardii* genome. We found that NatR-containing *S. boulardii* was the only microbe from mouse feces able to grow on YPD plates containing 50 µg/ml nourseothricin, 0.125 mg/ml Penicillin and 0.25 mg/ml streptomycin (data not shown). We then provided conventional mice with 1mg/ml penicillin and 2mg/ml streptomycin in their drinking water for four days, after which antibiotics treatment ceased and 10^8^ CFU of *S. boulardii* was delivered once per day for three days. Under this regimen (Treatment 1, Figure 4b), we found that *S. boulardii* titers fell to undetectable levels within 48 hours of the last gavage (Figure 4D). On the other hand, when antibiotic treatment was maintained over the course of the experiment (Treatment 2, Figure 4c), the titer and residence time of *S. boulardii* both increased, enabling *S. boulardii* to remain present in the gut one additional day after the last gavage (Figure 4D). These results are in line with previous *S. boulardii* colonization studies performed in mice colonized by a human microbiota^58^. Taken together, these experiments indicate that competitive microbial growth reduces the residence time of *S. boulardii* in the mouse gut. In these contexts, *S. boulardii*-mediated drug delivery will only occur during and for several days after probiotic administration, thereby minimizing dosage outside of desired therapeutic timescales.

**Figure 4.**
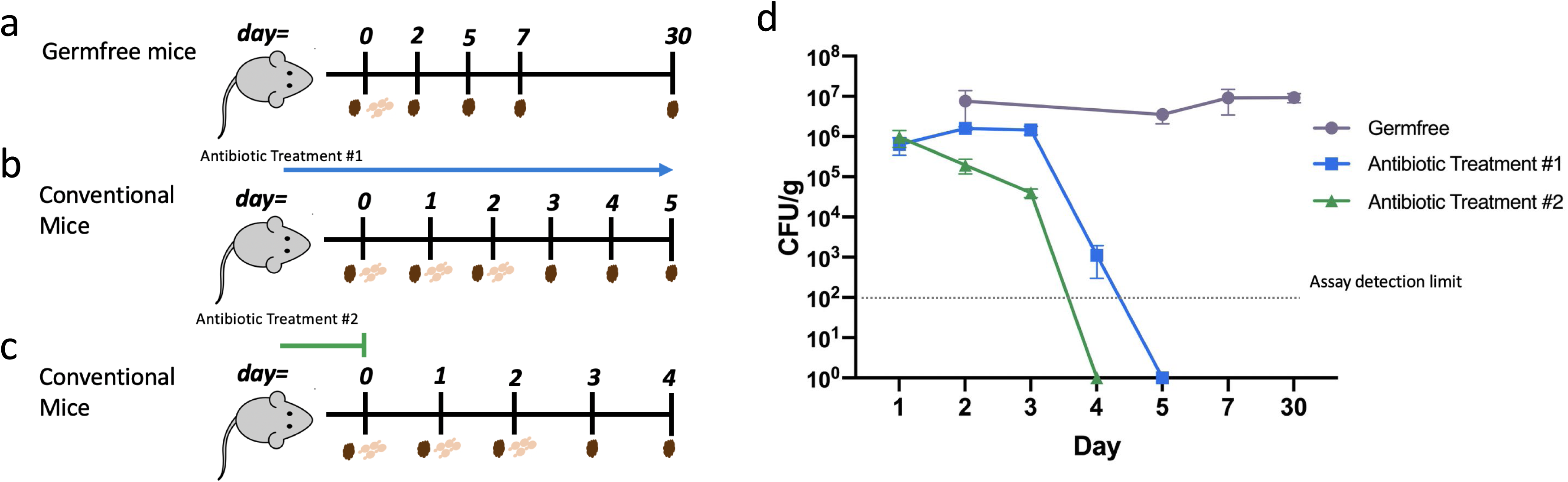
Microbial competition reduces *S. boulardii* residence time in mice. **a.** Germfree mouse model gavaged with *S. boulardii* on day 0. **b** Conventional mouse model treated with 1mg/ml penicillin and 2mg/ml streptomycin in the drinking water for 4 days prior to 3 *S.b::NatR* treatments every 24 hours (antibiotics were replenished every 4 days.) **c**. Conventional mouse model treated with 1mg/ml penicillin and 2mg/ml streptomycin in the drinking water for 4 days prior to 3 *S.b::NatR* treatments every 24 hours (antibiotic treatment was stopped on day 0.) **d.** Fecal carriage of *S.b* in the three mice models. Fecal samples were collected as shown in the timelines and plated on YPD media for the gnotobiotic model and YPD containing antibiotics for the conventional mice models. The dashed line shows the detection limit of our assay. Error bars indicate the standard deviation observed among 4 germ free mice, 3 conventional mice treated with antibiotic treatment #1, and 3 conventional mice treated with antibiotic treatment #2. In all mouse models, feces were collected prior to Sb gavage during days when both occurred.

## Conclusion

The gut and the microbes that live within it play a major role in host health, and as such are promising targets for manipulation via small molecule or biologic drugs. Unfortunately, delivering drugs to this environment remains challenging due to the digestive action of the host, especially for protein-based therapies. Due to its probiotic properties and similarity to the biomanufacturing workhorse organism *S. cerevisiae*, we characterized the ability of *S. boulardii* to be engineered to secrete biomolecules in the mammalian gut. We first characterized the effect of different plasmid-based and genomic sequences on the expression of synthetic constructs. We next established efficient genome editing tools for this strain and revealed the effects of genomic integration site on synthetic construct performance. We further harnessed *S. boulardii*’s high rates of homologous recombination to combinatorially assemble pathways for vitamin (β-carotene) and drug (violacein) production. Finally, we measured *S. boulardii*’s residence time in germ-free and antibiotic-treated mice. Taken together, these results indicate that the decades of work developing *S. cerevisiae* as a platform for biomanufacturing and synthetic biology can be readily extended to *S. boulardii*. Coupled with the continued development of engineered commensal and probiotic bacteria, *in situ* biomanufacturing by gut-resident microbes has the potential to greatly reduce distributional complexity, improve site-specific delivery, and reduce production costs for a wide array of biomolecules.

## Supporting information

Supplementary File 1 - Plasmids

## Acknowledgments

The authors gratefully acknowledge the assistance of Dr. Susan Tonkonogy and Karen Flores in performing germ-free mouse experiments. The authors thank Dr. Yong-Su Jin for kindly providing *S. boulardii* knockout strains. They thank the lab of Dr. Albert Keung for assistance with flow cytometry measurements. They thank the lab of Dr. Milad Abolhasani for assistance with HPLC experiments. They also thank Dr. John E. Dueber for kindly sharing the MoClo-YTK Toolkit and raw data. pLM494 was a gift from Bernd Müller-Röber (Addgene #100539). LbCas12a was a gift from Roubos Lab (Addgene #101748). This work was supported by startup funds from North Carolina State University’s Chemical and Biomolecular Engineering Department. D.D. and I.S.A. were supported by NCSU CBE startup funds. I.S.A. was also supported by the Ministry of Higher Education - Oman.

## Methods

### Strains and Culture Media

*Escherichia coli* Top10, NEB 5α and NEB 10β were used for plasmid construction and maintenance. *E. coli* cells were grown in lysogeny broth (LB) (5g/L yeast extract, 10 g/L tryptone, 10 g/L NaCl) at 37 °C supplemented with ampicillin (100 µg/mL), kanamycin (50 µg/mL) or chloramphenicol (50 µg/mL). *Saccharomyces boulardii* ATCC-MYA796 was used for antifungal marker and origin of replication characterization. *Saccharomyces boulardii* Δ*URA3,* provided by Yong-Su Jin, was used for auxotrophic marker, origin of replication, promoter, and terminator characterization, as well as genome editing and metabolic pathway assembly. *Saccharomyces boulardii* Δ*TRP1* and Δ*HIS3* strains from the Jin Lab were used for auxotrophic marker and origin of replication characterization. Promoter and terminator characterization experiments were conducted in synthetic media containing 0.67% (w/v) Yeast Nitrogen Base Without Amino Acids (Sigma-Aldrich), 2% Glucose (Fisher Scientific), 1.92g/L Yeast Synthetic Media Drop-Out Mix without Uracil (Sigma-Aldrich). For auxotrophic marker characterization, synthetic media with 0.67% (w/v) Yeast Nitrogen Base Without Amino Acids (Sigma-Aldrich), 2% Glucose (Fisher Scientific), 1.92% Yeast Synthetic Media Drop-Out Mix without the appropriate nutrient (histidine, tryptophan, or uracil) (Sigma-Aldrich) was used. Antifungal marker characterization experiments were conducted in yeast extract-peptone-dextrose medium (50 g/L YPD broth (Sigma-Aldrich)) supplemented with various concentrations (50 µg/mL, 100 µg/mL and 200 µg/mL) of geneticin (G418), zeocin, nourseothricin and hygromycin.

### Plasmid Construction & Cloning

All plasmids used in this work are listed in Supplementary File S1. A synthetic toolkit (MoClo-YTK) containing yeast parts were gifts from the Dueber Lab (Addgene #:1000000061). Characterization plasmids consisted of 8 parts: 2 connectors, promoter, coding sequence, terminator, yeast marker, yeast origin, and *E. coli* marker and origin. Each of these plasmids were assembled according to Deuber lab YTK protocols via Golden Gate cloning^59^. The Golden Gate reaction mixture, for all plasmids except those containing β-carotene or violacein pathways, was prepared as follows: 0.5 µL of 40 nM of each YTK plasmid (20 fmol), 0.5 µL T7 Ligase (NEB) or 0.5 µL T4 Ligase (NEB), 1 µL T4 Ligase Buffer (NEB) and 0.5 µL BsaI (10000 U/mL, NEB) and water to bring the final volume to 10 µL. The Golden Gate assembly protocol was performed on a thermocycler with the following program: 30 cycles of digestion (42 °C for 2 minutes) and ligation (16 °C for 5 minutes), followed by a final digestion (60 °C for 10 minutes) and heat inactivation (80 °C for 10 minutes).

### Yeast Competent Cells and Transformations

We optimized the yeast competent cell preparation and transformation protocol from Gietz et. al for *Saccharomyces boulardii*^60, 61^. For the characterization experiments, we used frozen competent cells, whereas for the genome editing experiments we used unfrozen competent cells prepared on the same day^60, 61^. To prepare competent cells, yeast colonies were inoculated into 1mL YPD and incubated in a shaking incubator overnight at 37 °C, 250 rpm. This culture was diluted into fresh 50 mL YPD (approximately OD600=0.25), and grown for 6-8 hours until OD600=1. The culture was centrifuged at 3000 xg for 5 min at room temperature to pellet the cells. After decanting the media, the pellet was resuspended in 25 mL of sterile water, and then pelleted at 3000 xg for 5 minutes. This pellet was resuspended in 1 mL of sterile water. The cells were then pelleted down at 10000 xg for 1 minute and water was discarded. The cells were then resuspended in 500 µL of frozen competent cell solution consisting of 2.5% (v/v) glycerol (Sigma-Aldrich) and 5% (v/v) DMSO (Thermo-Fisher). 50 µL of this suspension was aliquoted into 1.5 mL microcentrifuge tubes and stored in -80 °C. For transformations, frozen cells were thawed at 37 °C for 1 minute. Then, the cells were pelleted by centrifugation at 3000 xg for 2 minutes, after which the supernatant was removed. Then, solutions were added in the following order; 260 µL 50% PEG3350 (Fisher Scientific), 36 µL 1M Lithium Acetate (Sigma-Aldrich), 50 µL of 2 mg/ml single-stranded salmon sperm DNA ((Invitrogen™, 15632011) 10 mg/ml double-stranded salmon sperm DNA was diluted to 2 mg/ml and heat treated at 95 °C for 5 minutes to denature the double-stranded DNA), 0.1-10 µg DNA and water to bring the final volume to 360 µL. The pellet was resuspended in the transformation mix by gently mixing with a pipette tip. This transformation mix was incubated at 42 °C for one hour. This mixture was then centrifuged at 3000 xg for 1 min and the supernatant was discarded. The cell pellet was resuspended in 1 mL YPD by gently pipetting up and down and this tube was incubated at 37 °C for one hour. Then, the cell suspension was centrifuged for one minute at 3000 xg, resuspended in 25 µL sterile water, and plated on an appropriate growth media.

### S. boulardii Colony PCR

Since *S. boulardii* has a thicker cell wall than *S. cerevisia*e, we developed a new method to perform colony PCR on *S. boulardii*. Colonies were picked into 25 µL 20 mM sodium hydroxide and incubated at 98 °C for 80 minutes. Then we centrifuged these reactions at max speed (Mini Microcentrifuge Sargent-Welch) for 10 minutes and started a 25µL PCR reaction with 2µL of this reaction as a template. Otherwise, PCR reactions were performed according to the supplier’s instructions.

### Flow Cytometry

Constructs were inoculated from -80C freezer stocks into 750 µL of appropriate media in 96-deep-well plates (VWR International) and incubated at 37 °C for 36 hours with shaking at 250 rpm. After 36 hours, the cultures were diluted 1:100 in 200 µL of fresh media in 96-well-plates (Costar) and were grown for ∼8-10 hours at 37 °C. Once the cultures reached OD600 values between 0.1-0.4, flow cytometry was performed using a MACSQuant VYB (Miltenyi Biotec). Cells containing empty vector were used as a control. The voltage of the channels were adjusted in order to tune the gain of the detectors. The voltage values applied to FSC, SSC, Y1 (PE-A, excitation:561, emission: 586/15 nm), V1 (CFP-VioBlue, excitation: 405 nm, emission: 450/50 nm) and B1 (GFP_FITC, excitation: 488 nm, emission: 425/50 nm) channels/filters were 326V, 290V, 396V, 324V and 265V respectively. Flow cytometry data analysis was conducted on the FlowJo program (FlowJo LLC). The mean fluorescence values obtained from the flow cytometry for each transcriptional construct was then normalized to background fluorescence (i.e. the cells with an empty vector).

### Mouse Experiments

All mouse experiments were approved by the NC State University Institutional Animal Care and Use Committee (IACUC).

#### Germfree Mice

6-10 week old male and female germ free C57BL/6 mice were born at the NCSU Gnotobiotic Animal Core and were mono-colonized by *S. boulardii*. 2 mice (males) were gavaged with 10^6^ colony forming units in 100 µL PBS, 8 mice (6 females and 2 males) were gavaged with 10^7^ and 4 mice (2 females and 2 males) were gavaged with 10^8^ *S. boulardii* cells on day 1 and fecal samples were collected on days 2, 5 and 7. Fecal samples from mice housing the 10^8^ inoculum were collected on day 30.

#### Conventional Mice

Six-week old female C57BL/6J mice were obtained from Jackson Labs and hosted at the NCSU Biological Resources Facility (BRF) for 3-4 days before experiments. Mice were housed in groups of three and their cages were changed before treatment with antibiotics and before treatment with *S. boulardii* or water. All mice were treated with 1mg/ml penicillin and 2mg/ml streptomycin in the drinking water for 4 days starting on day -3. On day 1, drinking water was either replenished with antibiotic solution or changed to fresh water. Mice were then gavaged with 10^8^ CFU *S. boulardii* or water once every 24 hours for 3 days. Fecal samples were collected every 24 hours from day 1 to day 5.

#### Stool Cultures

1-2 pieces of stool were collected in weighed 1.5 ml centrifuge tubes and then weighed again to determine fecal mass. Fecal matter was then resuspended in 1 mL PBS per 100 mg feces. Fecal suspensions from mono-colonized mice were placed on solid YPD media, while fecal suspensions from conventional mice treated with antibiotics were plated on YPD media containing 50 µg/ml nourseothricin and 0.25 µg/ml streptomycin. Plates were sealed with parafilm and incubated at 37 °C for 2-3 days.

### Genome Editing

Integration plasmids contained *URA3* to aid selection and contained homology arms between 350 to 900 base-pairs in length as shown in Supplementary Figure S7. For DNA homologous recombination without DSB (**dsDNA integration**), 2µg of linear donor DNA was transformed to 10^7^ *S. boulardii* (*S. boulardii ΔURA3*) competent cells prepared on the same day. For **SpCas9**, 2µg of Linear donor DNA and 1.5µg of gRNA plasmids were transformed to 10^7^ freshly prepared chemical competent *S. boulardii ΔURA3* cells containing SpCas9 plasmid. For **LbCas12a**, 2µg of Linear donor DNA and 1.5µg of gRNA plasmids were transformed to 10^7^ freshly prepared chemical competent *S. boulardii ΔURA3* cells LbCas12a plasmid.

Overnight cultures of *S. boulardii ΔURA3* were grown in YPD only for **Homologous Recombination,** *S. boulardii ΔURA3* containing the **SpCas9** plasmid were grown in YPD + 100 µg/mL nourseothricin, and *S. boulardii ΔURA3* containing the **LbCas12a** plasmid were grown in YPD + 200 µg/mL G418 were subinoculated to prewarmed YPD media with or without antifungals at an OD600 of 0.25 and incubated at 37 °C with shaking at 250 rpm until cultures reached an OD600 of 0.9-1.0. Cultures were then centrifuged at 3000 xg for 5 min. Cells were then washed twice with 0.5 and 0.1 volumes of sterile DI water at room temperature at 3000xg for 5 min. The cells then were transformed according to the protocol described above, with the exception that heat shocked cells were spun down, resuspended in YPD, and incubated for 3 hours at 37 °C. Transformed cells were then spread on Yeast Complete Synthetic Media (CSM) without uracil plates with or without antibiotics. Plates were incubated at 37 °C for 2-4 days. Integration was verified with colony PCR.

### Combinatorial Pathway Assembly

Prior to library assembly, promoter, gene, and terminator part plasmids were assembled into ‘entry vector’ plasmids by BsmBI assembly. These plasmids had BsaI overhangs and follow the ‘Yeast golden gate toolkit’ standard [deuber gg]. Backbones were assembled for subcloning of the transcriptional units. These backbones housed the appropriate homology arms for later final assembly of transcriptional units into a full pathway. BsaI assembly of transcriptional units into the matching backbone was completed by combining 10 fmol of each promoter part plasmid (to a total of 100 fmol) with 100 fmol of gene, terminator, and backbone plasmids. Reaction mixtures included 0.5 µL of BsaI-HF v2, 0.5 µL of T7 DNA Ligase, and 1 µL of T4 DNA ligase buffer (NEB), and underwent 50 cycles of golden gate assembly (2 minutes at 42 °C, 5 minutes at 16 °C), before a single 30 minute final digestion at 60 °C, and 10 minutes of heat inactivation at 80 °C. Subsequently, we have noticed that final digestion steps of 5 minutes at 42 °C are more optimal for BsaI assembly^59^. Transcriptional units were amplified directly from the golden gate mixture using 1 µL of reaction mixture as template in 50 µL Q5 Hot-Start Master Mix reactions (NEB). PCR reactions were performed according to manufacturer’s instructions and cleaned up using clean and concentrate kits (D4004-Zymo Research). 600 nM of each amplified transcriptional unit library was transformed using the lithium acetate protocol described above. Cells were recovered for 1 hour in YPD prior to dividing the culture into 5 aliquots of 200 µL and plating on yeast nitrogen base plus amino acids without uracil (Sigma).

### Pathway Sequencing

Random colonies were collected in 1.5 centrifuge tubes and resuspended in 1 ml of 10 mg/ml Lysing Enzymes from *Trichoderma harzianum* (L1412 Sigma). The cells were incubated in the lysing enzyme at 37 °C for 3 hours and then the ZymoPURE Plasmid Miniprep (D4208T) was used to miniprep the plasmids. The Lysing Enzymes from Trichoderma harzianum help digest *S. boulardii*’s wall and produce protoplasts that are easy to miniprep. 2µl of the miniprep product was then used as a template to amplify the assembled region with primers complementary to the pathway genes or terminators, as the ordering of these was common to all constructs. The PCR product was then sent for Sanger sequencing (Genewiz, NJ,USA) to verify the promoters assigned for each gene.

### Antifungal Minimum Inhibitory Concentration tests in *S. boulardii*

3 biological replicates of *S. boulardii* (ATCC-MYA796) were grown overnight in YPD media. Overnight cultures were sub-inoculated into YPD media with different antifungal concentrations ranging from 0 µg/ml to 200 µg/ml in a 96 well plate at starting OD600 0.02 and grown for 36 hours in a plate reader (BioTek Synergy™ H1, Shake Mode: Double Orbital, Orbital Frequency: continuous shake 365 cpm, Interval: 10 minutes). Geneticin (G418), Nourseothricin, Zeocin, Hygromycin B were used in this study.

### Extraction of β-carotene from *S. boulardii* cultures

3 replicates of each β-carotene producing *S. boulardii* strain were inoculated into 1 mL CSM without uracil (2.5% glucose) and and grown overnight in a shaking incubator at 250 rpm at 37 °C. Overnight cultures were then subinoculated (1:100) into 30 mL CSM without uracil (2.5% glucose) and grown for 3 days in a rotary incubator at 250 rpm at 37 °C. After 3 days, 2*10^9^ cells were collected by centrifuging at 4 °C at 5000g for 5 minutes. The media was discarded and the cells were washed with 10 mL water and were centrifuged at 4 °C at 5000g for another 5 minutes. After the water was discarded, the cells were resuspended in 500 µL water and transferred into ZR Bashing Bead Tubes (Zymo Research, CA). The ZR Bashing tubes were centrifuged for 1 minute at 8000g to discard the water. 500 µL acetone was then added into the bashing tubes. The tubes containing the cells, acetone, and lysis beads then transferred to a homogenizer (TissueLyser, Qiagen, Germany) for physical lysis (Lysis duration: 10 minutes, frequency: 30/s). The cells and the lysates then centrifuged at 16000g for 5 minutes at 4 °C. The supernatants (acetone lysate) were transferred to 1.5 mL black microcentrifuge tubes. Fresh acetone was then added to the bashing tubes, and lysis steps were repeated 2 more times, at which point the cell pellets were completely white. β-carotene extracts were dried by speed vac (aqueous solvent, 30 °C for 2 hours). Dried β-carotene pellets were reconstituted in 500 µL acetone and stored at -20 °C in the dark.

### Detection of β-carotene by HPLC

5µL of each sample was processed in an Agilent 1260 Infinity high performance liquid-chromatography (HPLC) system (Agilent Technologies, CA, USA). 0.002 mM naphthalene solution was used as internal control and pure β-carotene (Thermo Fisher, Waltham, MA) at varying concentrations (mg/mL) 0.00625, 0.0125, 0.025, 0.05, 0.0625, 0.125, 0.25, 0.5, 0.625, 1.25, 2.5, 5) as external control. The channels used for internal control and β-carotene detection were 275 nm and 453 nm, respectively. The column used in the system was an Agilent HC-C28 Reversed Phase column (4.6 x 250 mm, 50 µL) (Agilent Technologies, CA, USA). The composition of the mobile phase was acetonitrile:methanol:isopropanol (v/v/v: 85/10/5) The flow rate for the mobile phase was 3 mL/min.

---

**Figure S1.**
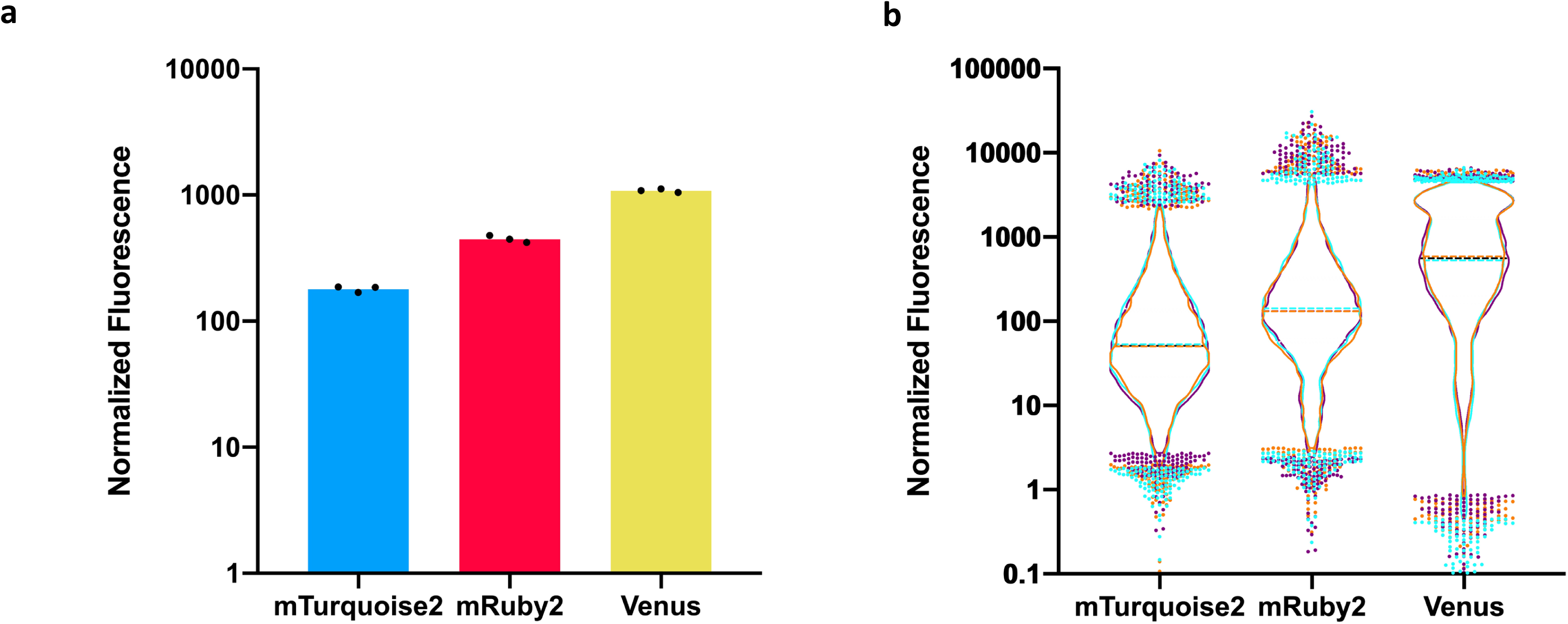
Characterization of Fluorescent Genes. **a)** 3 fluorescent genes (*mTurquoise2, mRuby2* and *Venus*) were cloned behind *TDH3* promoter and in front of *TDH1* terminator. The height of the bars represent the mean fluorescent value of 3 biological replicates (n=3). The black dots show the normalized mean of each biological replicate. **b)** Violin plots of flow cytometry data for 3 fluorescent proteins. Cyan, orange and purple represent first, second and third biological replicates, respectively. The dots represent the highest and lowest 2 percent of the cell population. Each flow cytometry run consists of 10000 events.

**Figure S2.**
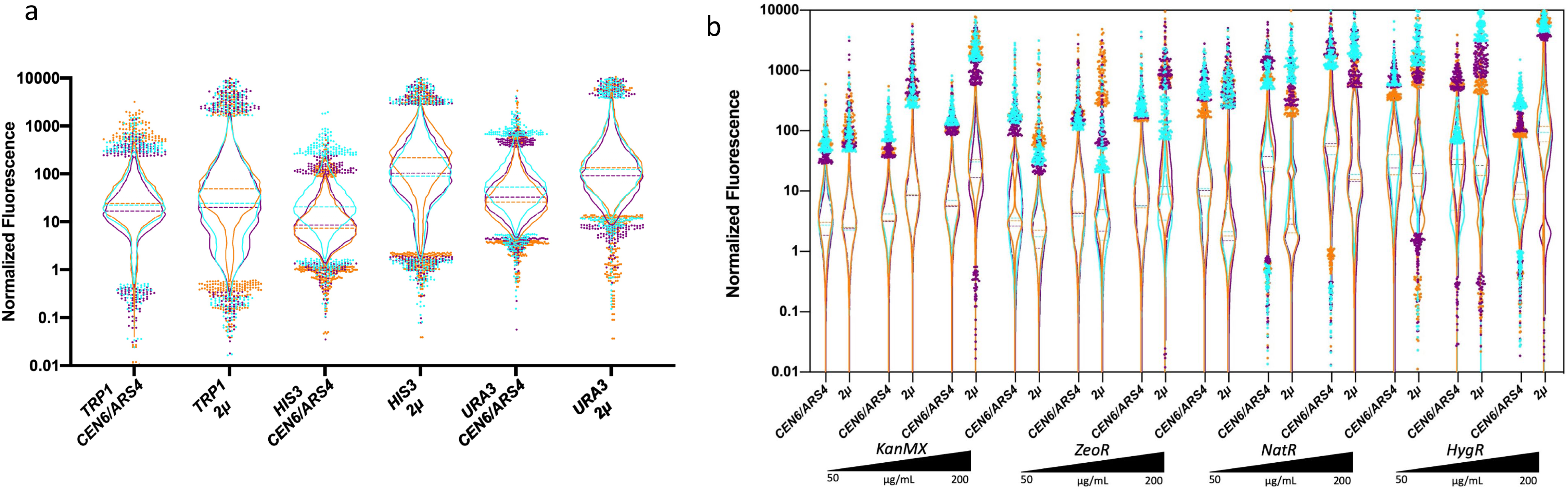
Characterization of selective markers and origins of replication. **a)** Violin plots of flow cytometry data for 3 auxotrophic markers and 2 origins of replication. **b)** Violin plots of flowcytometry data for 7 antifungal markers and 2 origins of replication tested in 3 different antifungal concentrations (50, 100, 200 μg/mL). 3 auxotrophic markers, 4 antifungal markers and 2 origins of replications were cloned behind *TDH3* promoter, mRuby2 fluorescent protein and *TDH1* terminator. Cyan, orange and purple represent first, second and third biological replicates, respectively. The dots represent the highest and lowest 2 percent of the cell population. Each flow cytometry run consists of 5000-10000 events.

**Figure S3.**
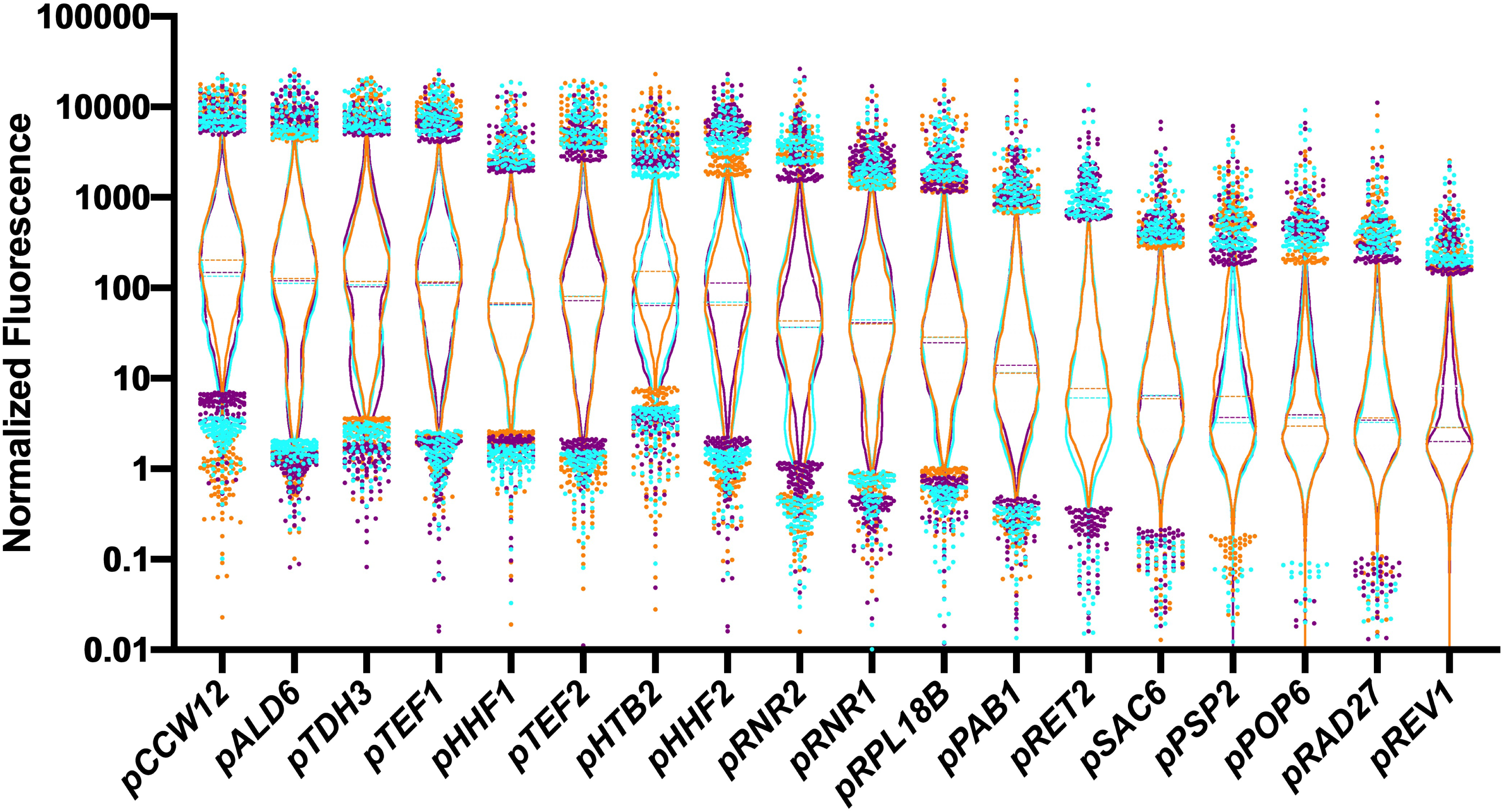
Characterization of Promoters. Violin plots of flowcytometry data for 18 constitutive promoters. 18 constitutive promoters were cloned in front of mRuby2 fluorescent protein and *tTDH1* terminator. Cyan, orange and purple represent first, second and third biological replicates, respectively. The dots represent the %2 percent of the cell population. Each flowcytometry run consists of 10000 events.

**Figure S4.**
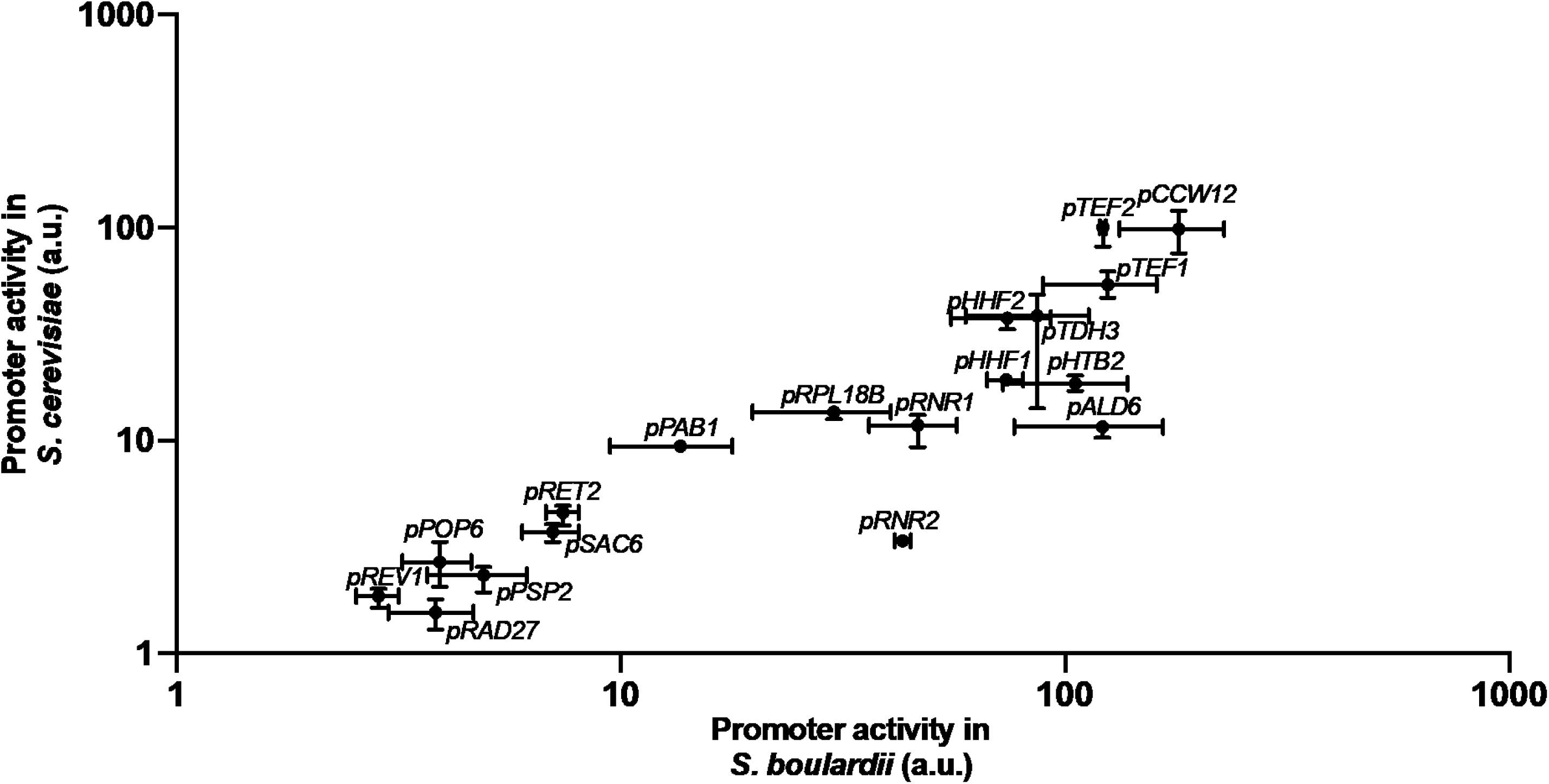
Comparison of constitutive promoter activity between *S. cerevisiae* and *S. boulardii*. Promoter activity of 18 constitutive promoters in *S. boulardii* (x-axis) and *S. cerevisiae* (y-axis) were plotted (R=0.81). In each synthetic construct, one constitutive promoter was cloned upstream of *mRuby2* gene. Activity of each promoter in both yeast species was determined by reading median fluorescence values produced by mRuby2 protein via flowcytometry. Dots correspond to the average median of 3 (*S. boulardii*) and 4 (*S. cerevisiae*) biological replicates and the bars correspond to maximum and minimum median value obtained by the replicates.

**Figure S5.**
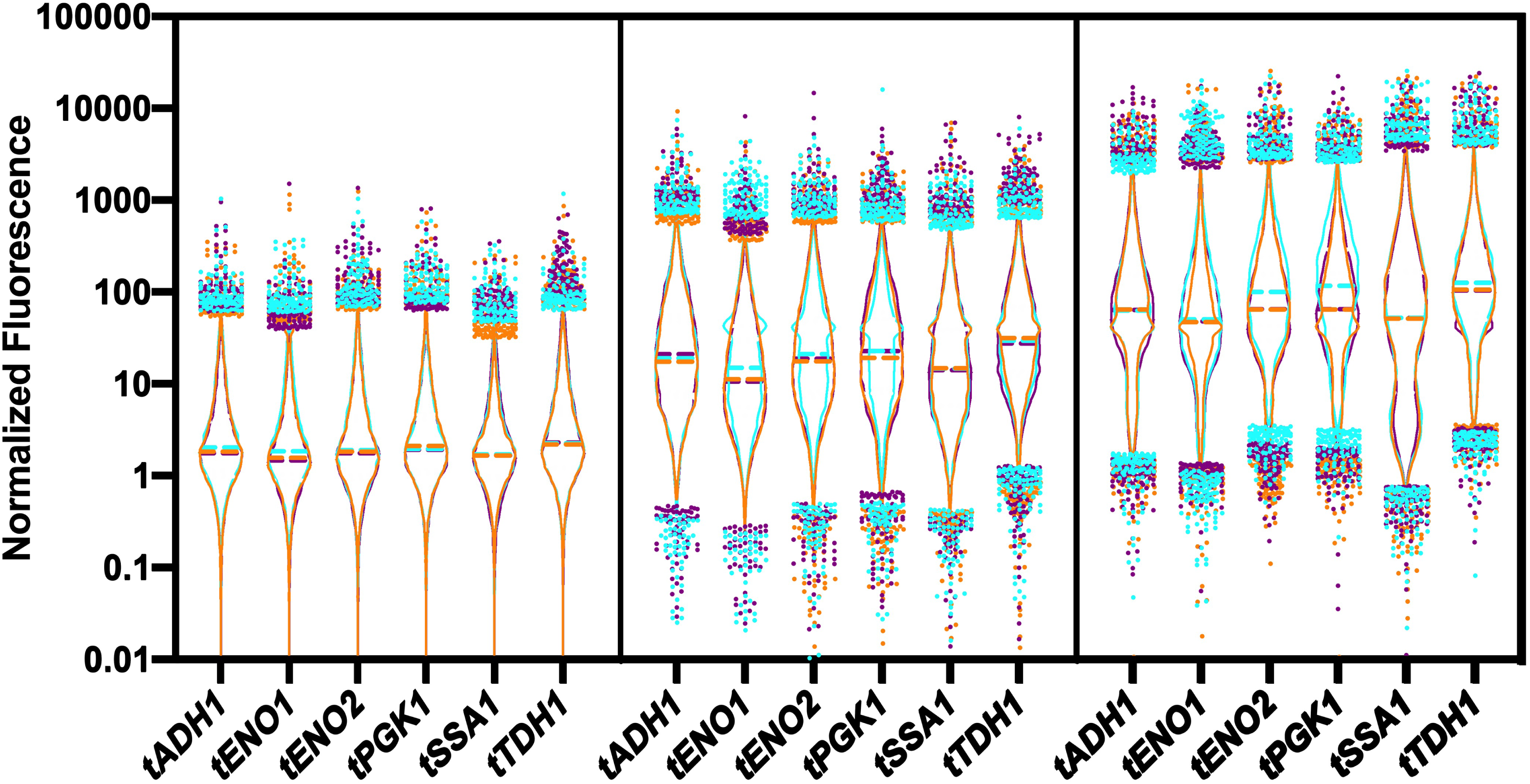
Characterization of Terminators. Violin plots of flowcytometry data for 5 terminators combined with 3 promoters. 5 terminators were cloned behind 3 promoters with different strength (*pTDH3*-strong-most right panel, *pRNR1*-medium-middle panel, *pREV1*-weak-most left panel) and mRuby2 fluorescent protein. Cyan, orange and purple represent first, second and third biological replicates, respectively. The dots represent the %2 percent of the cell population. Each flowcytometry run consists of 10000 events.

**Figure S6.**
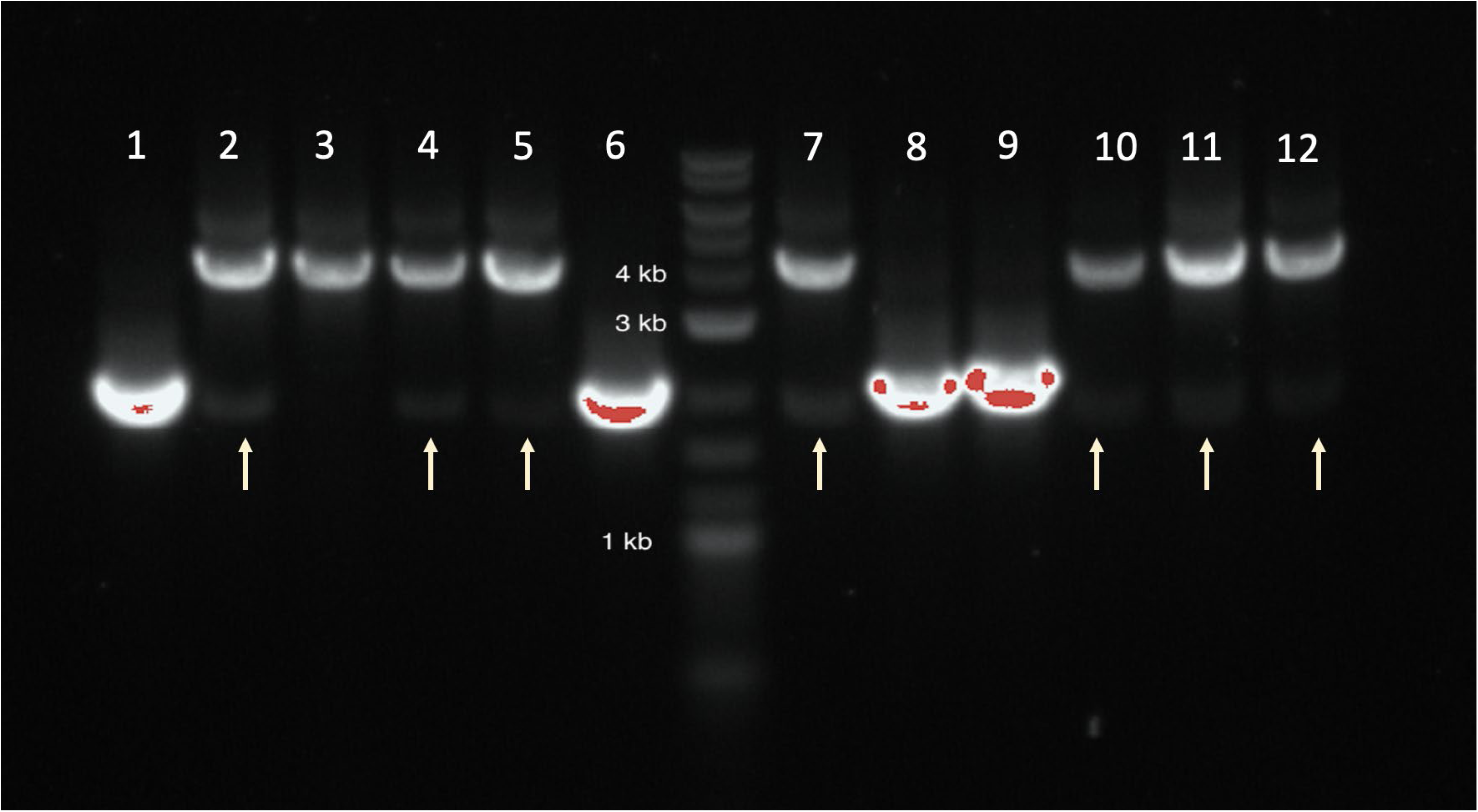
Colony PCR of *S. boulardii* genome editing using only homologous recombination. The gel image shows that when only donor DNA is transformed into *S. boulardii*, the cells keeps a copy of the original chromosomal locus sequence. Lanes 1-6 lanes show a PCR for correct integration into INT1. Lanes 7-12 show a PCR for correct integration into INT4. In both cases, a correct integration will yield a PCR product of 3.8 kb, while the native sequence yields a PCR product of 2.0 kb.

**Figure S7.**
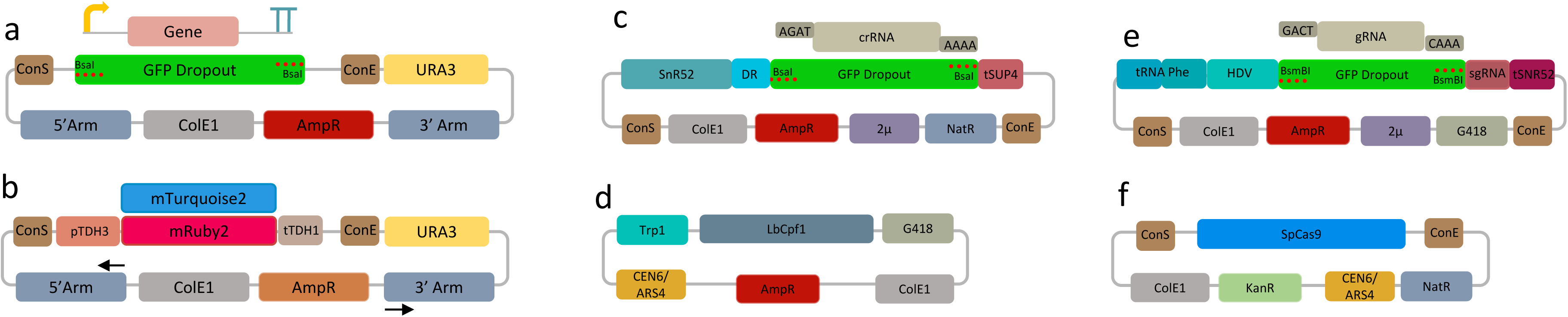
Genome editing plasmids. **a.** Vector for assembling the promoter, gene and terminator of interest. **b.** Example of constructed integration plasmid using cloning vector from a. **c.** Cloning vector with GFP dropout for constructing the CRISPR/LbCas12a crRNA **d.** CRISPR/LbCas12a expression vector. **e.** Cloning vector with GFP dropout for constructing the CRISPR/SpCas9 gRNA. **f.** CRISPR/SpCas9 expression vector. All GFP dropout vectors vectors are used in Golden Gate reactions with either BsaI or BsmBI enzymes.

**Figure S8.**
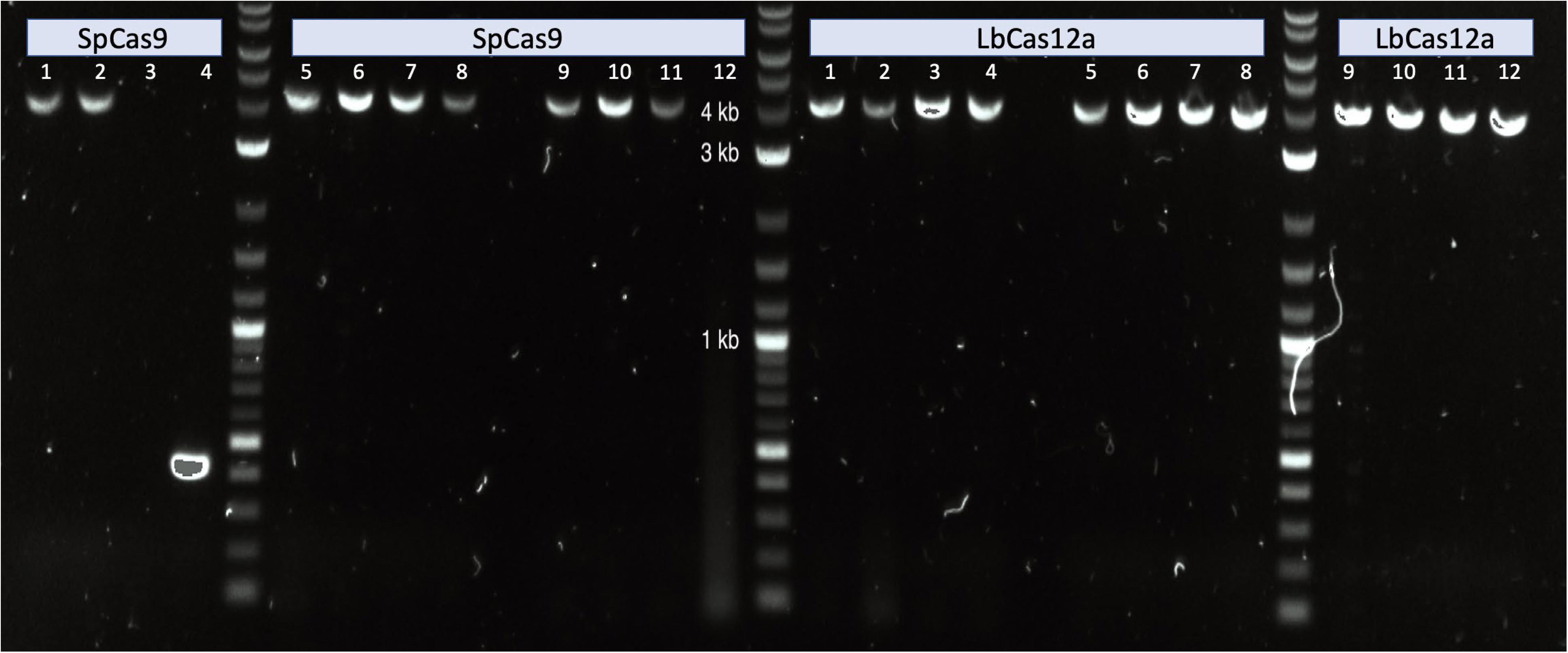
Colony PCR of *S. boulardii* genome editing using CRISPR/SpCas9 and CRISPR/LbCas12a. SpCas9 lanes 1-4 show a PCR screen for integration into INT1 (line 3 is failed PCR reaction and line 4 is unedited site), SpCas9 lanes 5-8 show a PCR screen for integration into INT4, and SpCas9 lanes 9-12 show a PCR screen for integration into INT5. LbCas12a lanes 1-4 show a PCR screen for integration into INT1, LbCas12a lanes 5-8 show a PCR screen for integration into INT4 and LbCas12a lanes 9-12 show a PCR screen for integration into INT5.

**Figure S9.**
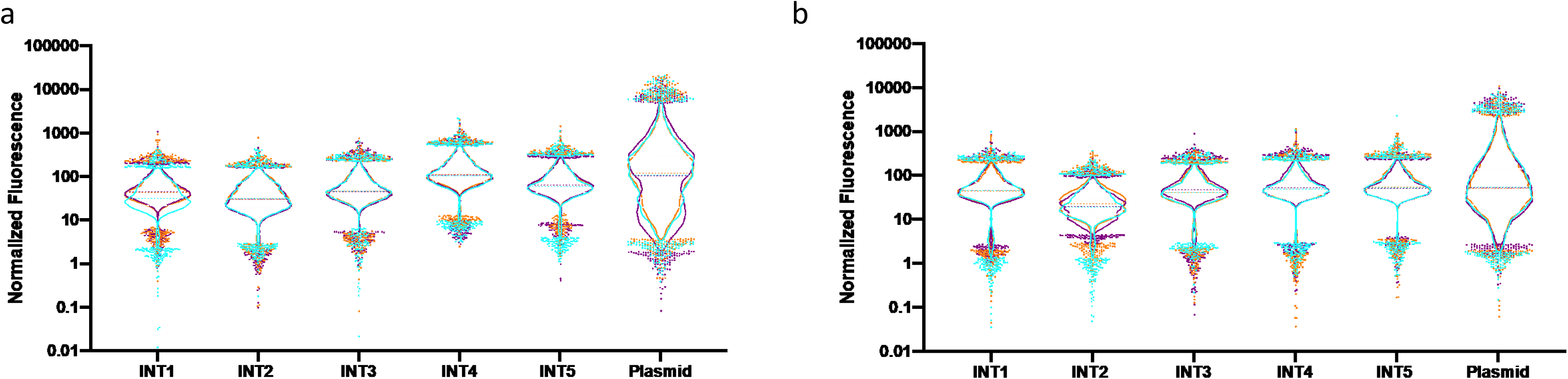
Effect of chromosomal locus on the expression of the fluorescent genes *mRuby2* and *mTurqouise2*. Violin plots of flowcytometry data for **a.** *mRuby2* expression in 5 different loci in the genome compared to the expression of the same construct in plasmid with *2µ.* b. *mTurqiouse2* expression in 5 different loci in the genome compared to the expression of the same construct in plasmid with *2µ* origin. constitutive promoters.

**Figure S10.**
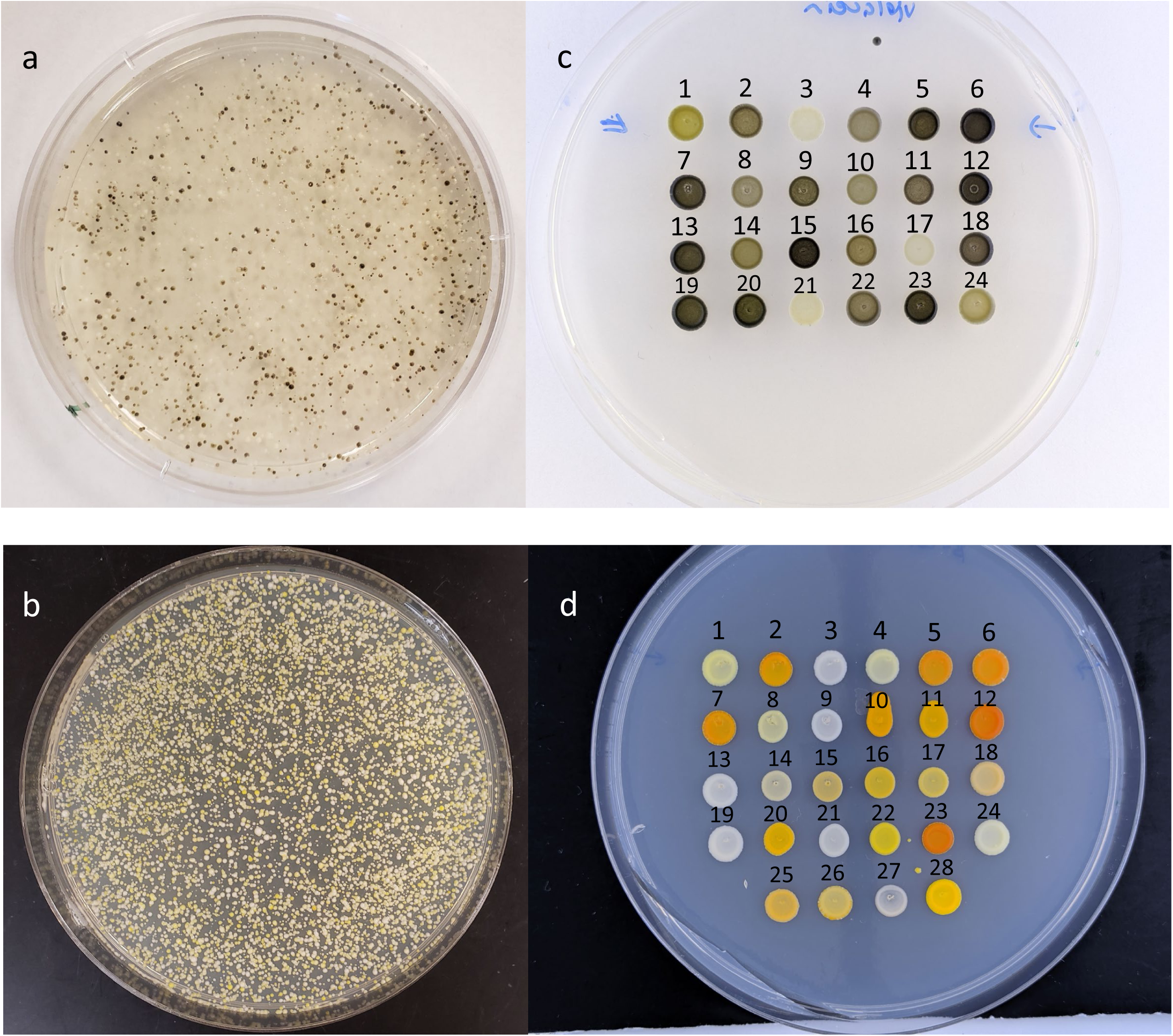
*in vivo* Combinatorial Pathway Assembly in *S. boulardii*. **a&b)** *S. boulardii* Transformation plates for violacein (a) and β-carotene (b) pathways. Violacein **(c)** and β-carotene (d) isolates picked from plates a &b.

**Figure S11.**
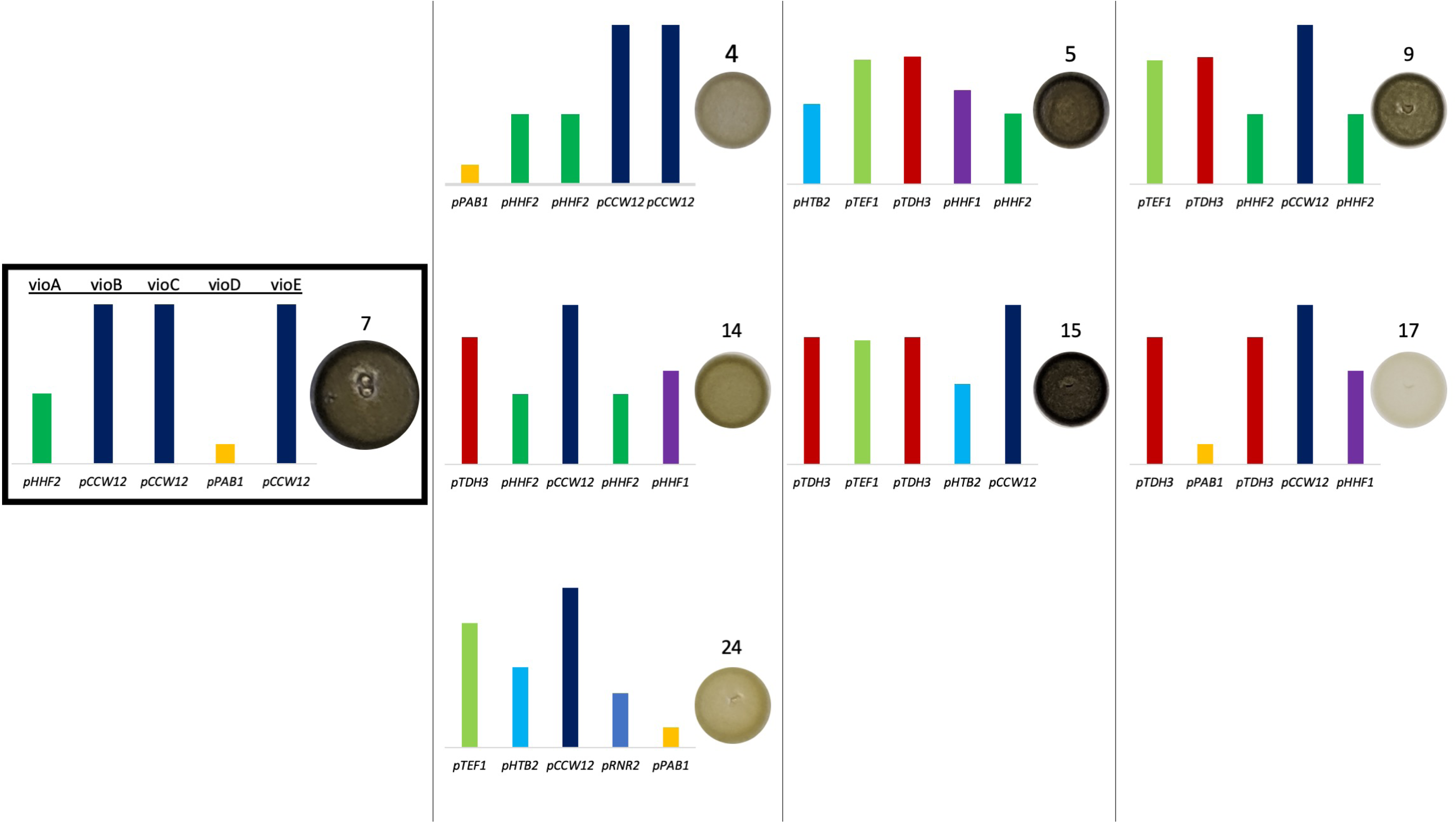
Combinatorial assembly of violacein pathway assembly in *S. boulardii*. Violacein pathway consisting of 5 genes (*vioA*, *vioB*, *vioC, vioD* and *vioE*) were assembled similar to β-carotene pathway. 8 out of 24 isolates (Supplementary Figure S8) were selected to proceed with sequencing based on stability of colony color after 3 serial passages. Sanger sequencing was done in a similar manner to the β-carotene isolates. Numbers on the colonies corresponds to the colony number from isolate plates on Figure S8c. The bar heights represent the normalized fluorescence values in linear scale for each promoter according to promoter characterization work (Figure 1a)

**Figure S12.**
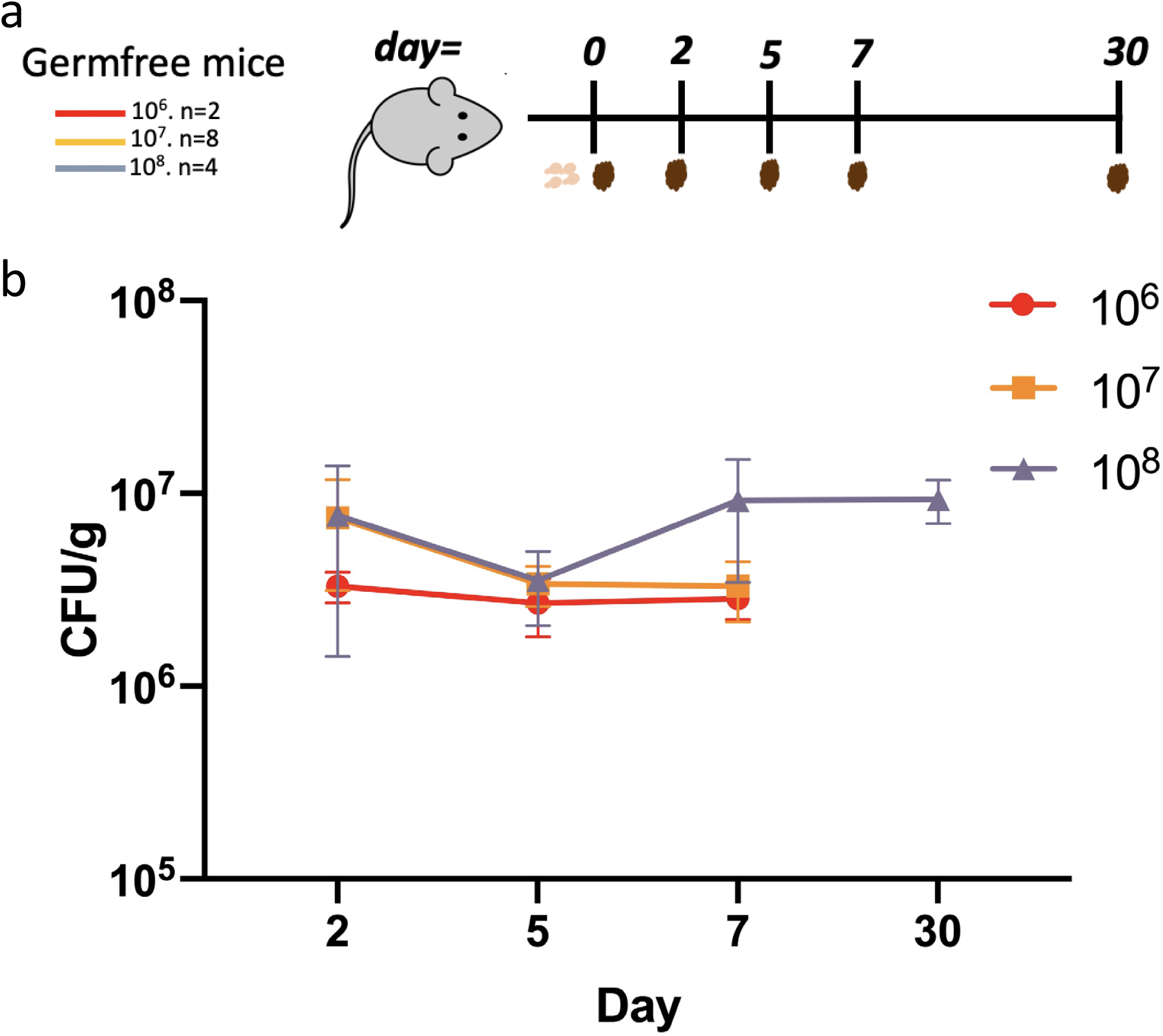
Dosage impacts the residence time of *S. boulardii* in germfree mice. **a**. germfree mice were gavaged with 3 different amounts of *S. boulardii* (10^6^, 10^7^ and 10^8^ cells) on day 0. **b**. Residence time of *S. boulardii* in germfree mice. Fecal samples were collected as shown in the timeline. Error bars indicate the standard deviation observed among 2 germ free mice for 10^6^ CFU treatment, 8 germ free mice for 10^7^ CFU treatment, and 4 germ free mice for 10^8^ CFU treatment In all mouse models. Feces were collected prior to Sb gavage during days when both occurred.

**Figure S13.**
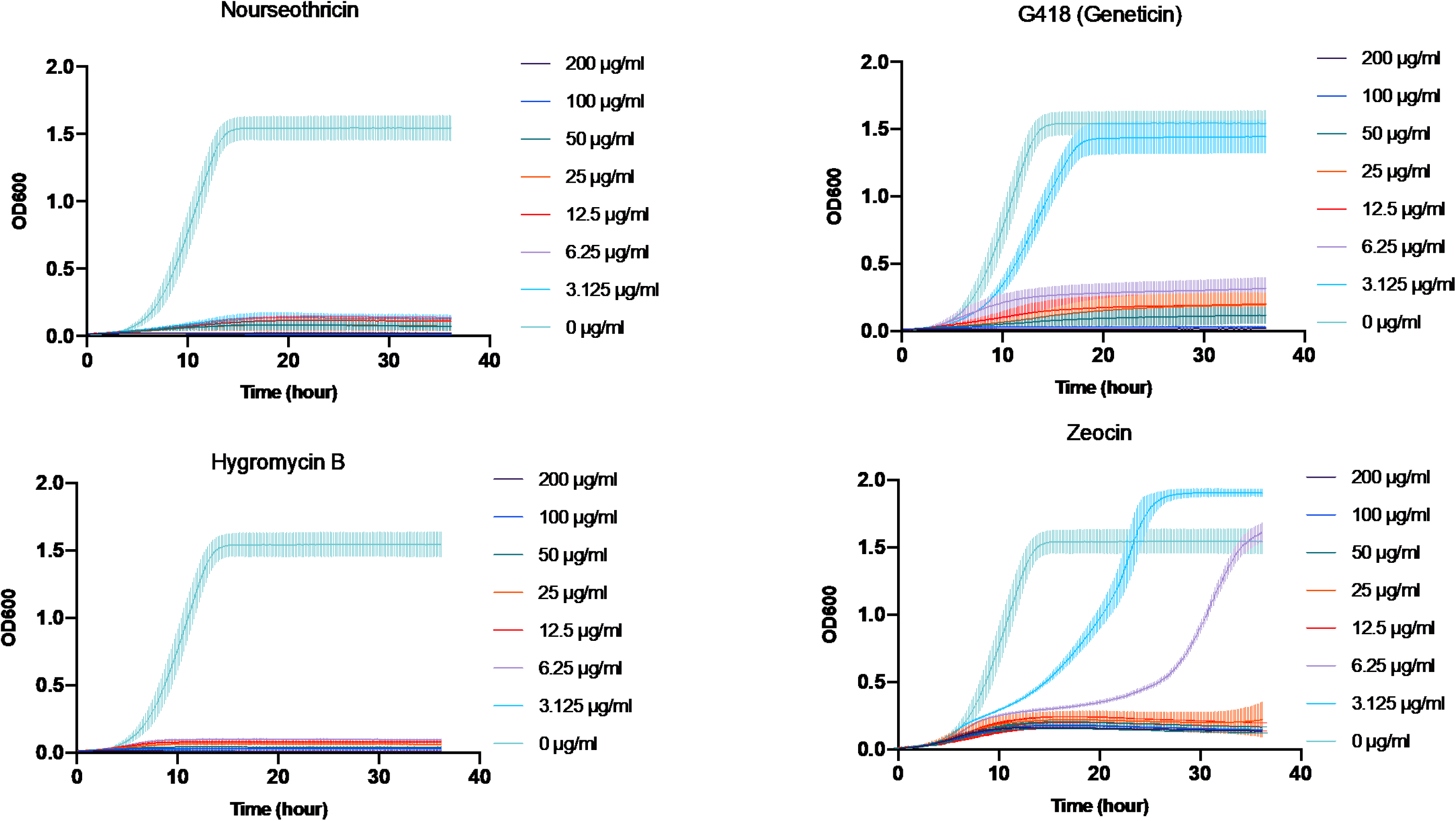
*S. boulardii* Minimum Inhibitory Concentration (MIC) Tests with Nourseothricin, G418 Geneticin, Hygromycin B and Zeocin. Error bars indicate the standard deviation observed among 3 biological replicates.

**Table S1.** MoClo kit promoters blasted against *S.boulardii’s* genome.

| **MoClo kit** | **Promoter** | **Score** | **Expect** | **Identities** | **Gaps** |
| --- | --- | --- | --- | --- | --- |
| YTK009 | *pTDH3* | 1218 bits(659) | 0 | 673/680(99%) | 0/680(0%) |
| YTK010 | *pCCW12* | 1234 bits (668) | 0 | 690/700 (98%) | 4/700 (0%) |
| YTK011 | *pPGK1* | 1282 bits(694) | 0 | 698/700(99%) | 0/700(0%) |
| YTK012 | *pHHF2* | 1197 bits(648) | 0 | 667/676(99%) | 1/676(0%) |
| YTK013 | *pTEF1* | 1271 bits(688) | 0 | 696/700(99%) | 0/700(0%) |
| YTK014 | *pTEF2* | 917 bits(496) | 0 | 534/550(97%) | 11/550(2%) |
| YTK015 | *pHHF1* | 1254 bits(679) | 0 | 694/701(99%) | 1/701(0%) |
| YTK016 | *pHTB2* | 1210 bits(655) | 0 | 683/696(98%) | 4/696(0%) |
| YTK017 | *pRPL18B* | 1236 bits(669) | 0 | 690/700(99%) | 2/700(0%) |
| YTK018 | *pALD6* | 1266 bits(685) | 0 | 695/700(99%) | 0/700(0%) |
| YTK019 | *pPAB1* | 1271 bits(688) | 0 | 697/701(99%) | 1/701(0%) |
| YTK020 | *pRET2* | 1288 bits(697) | 0 | 699/700(99%) | 0/700(0%) |
| YTK021 | *pRNR1* | 1266 bits(685) | 0 | 695/700(99%) | 0/700(0%) |
| YTK022 | *pSAC6* | 1266 bits(685) | 0 | 695/700(99%) | 0/700(0%) |
| YTK023 | *pRNR2* | 1251 bits(677) | 0 | 695/703(99%) | 4/703(0%) |
| YTK024 | *pPOP6* | 1168 bits(632) | 0 | 688/712(97%) | 15/712(2%) |
| YTK025 | *pRAD27* | 1184 bits(641) | 0 | 678/694(98%) | 9/694(1%) |
| YTK026 | *pPSP2* | 1236 bits(669) | 0 | 689/699(99%) | 0/699(0%) |
| YTK027 | *pREV1* | 1251 bits(677) | 0 | 693/700(99%) | 4/700(0%) |
| YTK028 | *pMFA1* | 1271 bits(688) | 0 | 697/701(99%) | 1/701(0%) |

**Table S2.**
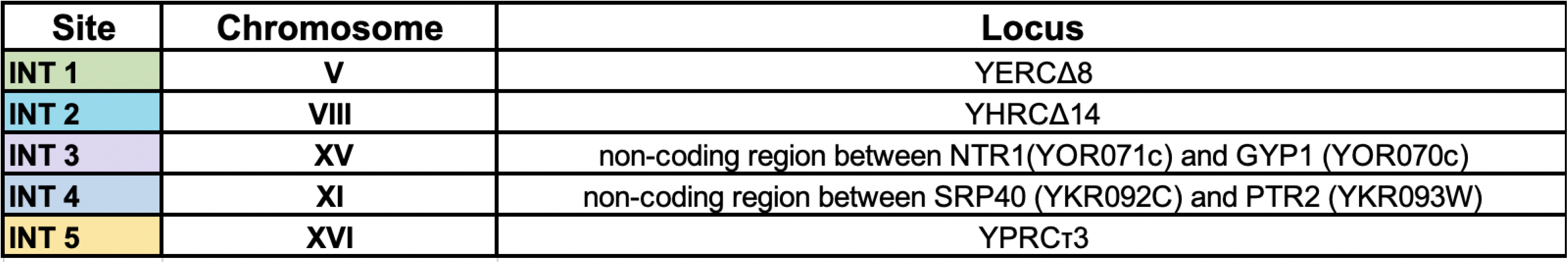
Integration locations (INT1-INT5) in *S. boulardii’s* genome.

**Table S3.**
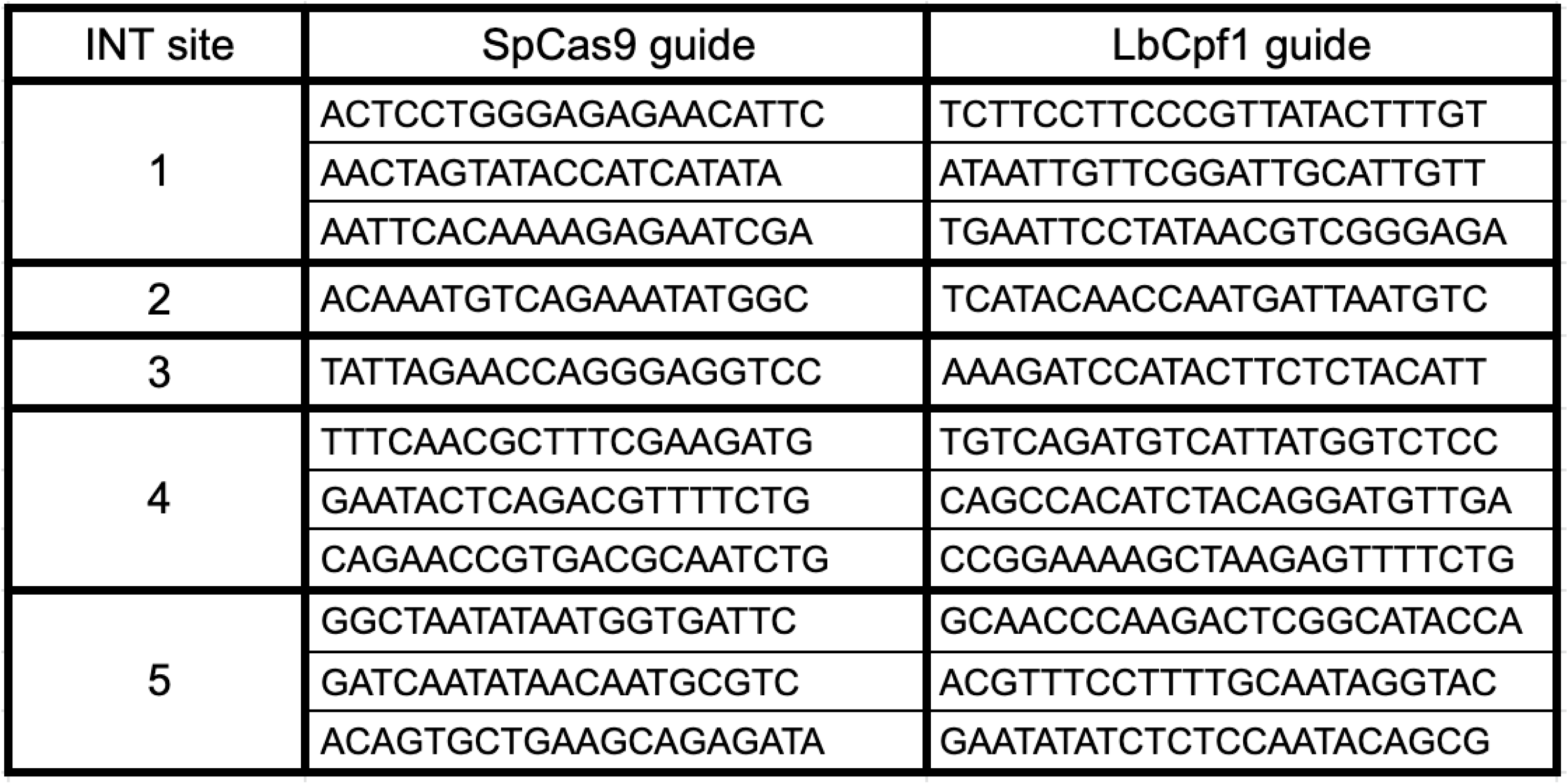
Guide RNA sequences used for genome editing assisted with the CRISPR nucleases SpCas9 or LbCas12a.

**Table S4.**
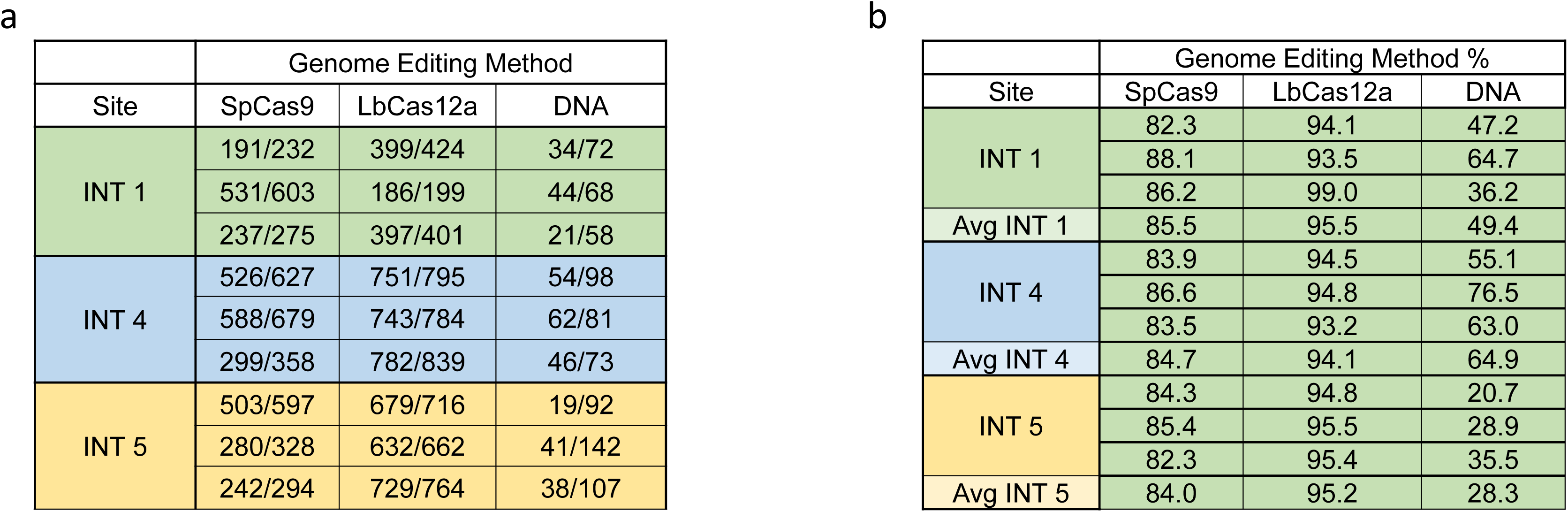
*S. boulardii* genome editing efficiencies using 3 different methods. **a.** Number of colonies screened for each editing method. Successfully edited cells are defined by the number of fluorescent (mTurquoise2) colonies divided by the total number of colonies on the plate **b.** The percentage of successfully edited cells in the 3 different locations using 3 different editing methods.

